# *Candidatus* Pannonibacter alzirae sp. nov. isolated from a septic immunocompromised patient in Brazil

**DOI:** 10.64898/2026.11.11.765890

**Authors:** Igor Vasconcelos Rocha, Beatriz Souza Toscano de Melo, João Luiz de Lemos Padilha Pitta, Carlos Alberto das Neves de Andrade, Reginaldo Gonçalves de Lima Neto, Ana Carolina Mota de Faria, Marilia Lavocat Nunes, Gabriel da Luz Wallau, Túlio de Lima Campos, Karina Lidianne Alcântara Saraiva, Maria Eduarda Bezerra de Andrade Carneiro, Marise Sobreira, Matheus Filgueira Bezerra

## Abstract

*Pannonibacter* species. are non-fermentative Gram-stain-negative bacteria, with rare reports of human infections primarily affecting immunocompromised patients. Here, we report the genomic and phenotypic characterization of a novel *Pannonibacter* species, isolated from a bacteremia episode in a pediatric patient with acute lymphoblastic leukemia in Brazil. Initial characterization of the isolate (here named CBAS 1226) by MALDI-TOF mass spectrometry and 16S ribosomal RNA gene sequencing identified it as *Pannonibacter* sp., most closely related to *P. phragmitetus*. However, comparison of its MALDI-TOF spectrum with a commercial reference library revealed compatibility with only two of the five main spectra profiles for *P. phragmitetus*. This discrepancy motivated comprehensive genomic, biochemical, morphological, and antimicrobial resistance profiling of CBAS 1226. Genomic analysis revealed that *Pannonibacter* sp. CBAS 1226 is phylogenetically distinct from *P. phragmitetus* and other known species, with ANI and dDDH values below species delineation thresholds, supporting its proposal as *Candidatus* Pannonibacter alzirae. It forms small, non-hemolytic, grayish-white colonies on blood agar, similar to its closest relatives *P. phragmitetus* and *P. indicus*. Phenotypically, it is distinguished by positive α– and β-glucosidase activity, along with the utilization of citrate, L-proline, and 2-ketoglutarate. It is negative for alkaline phosphatase and unable to utilize D-fructose, D-melibiose, esculin, or L-arginine. The isolate exhibited higher MICs for β-lactams, whereas carbapenems and fluoroquinolones showed low MICs. The isolate is capable of forming biofilms and tolerates alkaline conditions, a wide range of temperatures, and different oxygen concentrations. Altogether, this report emphasizes the relevance of additional proteomics– and genomics-based approaches to clarify unusual bacterial infections and describes a new *Pannonibacter species* with distinct features and the potential to cause opportunistic infections.

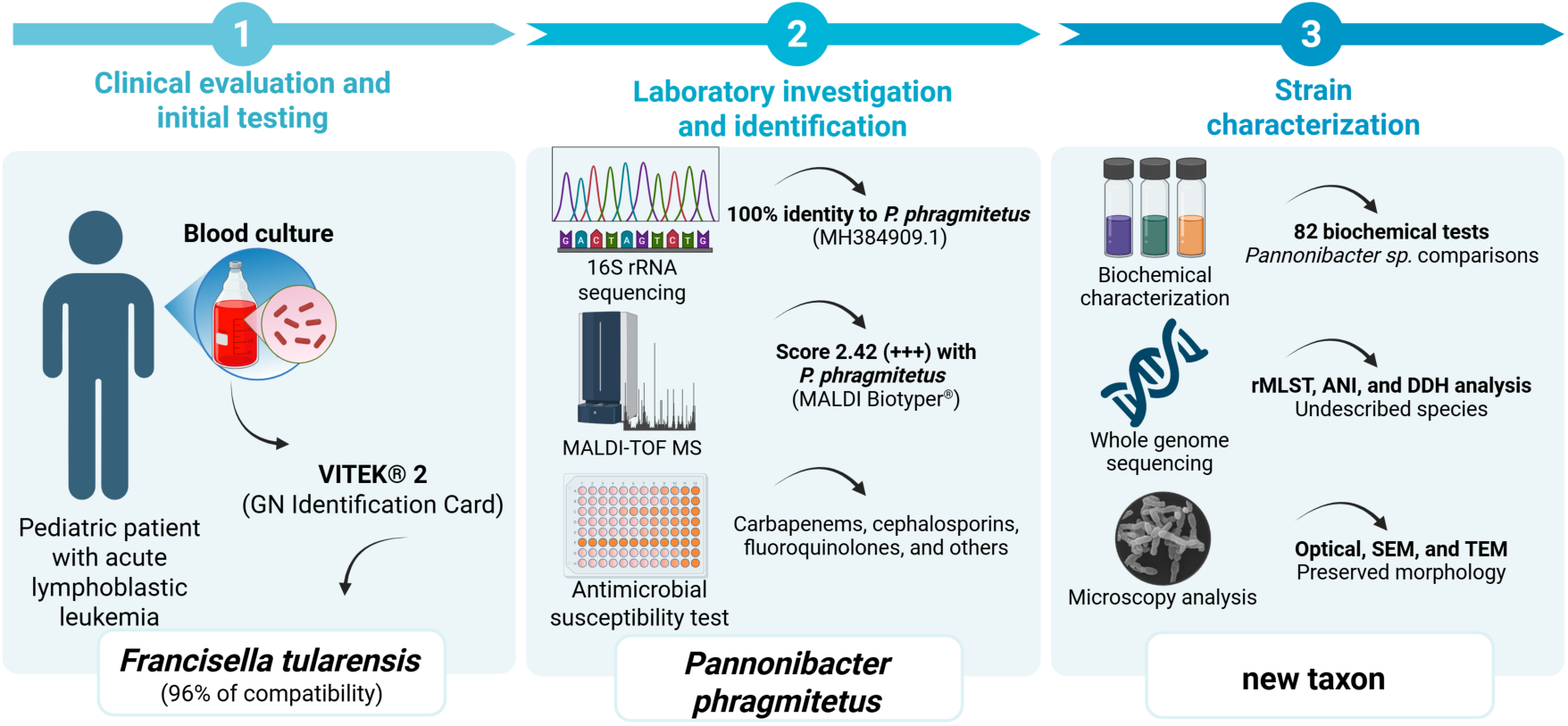
Graphical abstract.

## Introduction

*Pannonibacter* sp. consists of Gram-stain-negative, rod-shaped, facultatively anaerobic bacteria mostly known for their metabolic and environmental versatility, with potential applications in bioremediation processes [1, 2]. The genus currently comprises four species validly published under the International Code of Nomenclature of Prokaryotes (ICNP – *P. phragmitetus, P. indicus, P. carbonis and P. tanglangensis*), and one additional species awaiting valid publication (*P. anstelovis*, https://lpsn.dsmz.de/).

Members of the genus have predominantly been recovered from environmental habitats, including freshwater and soda lakes, hot springs, decomposing plant material, and coal mine-associated environments, indicating adaptation to diverse physicochemical conditions. *P. phragmitetus*, the type species of the genus, was originally isolated from decomposing reed rhizomes in a Hungarian soda lake and was characterized as an alkalitolerant bacterium [3]. *P. indicus* was subsequently isolated from a hot spring in India and demonstrated marked tolerance to arsenate, further illustrating the ecological and metabolic diversity of the genus [2]. This capacity for tolerance to environmental contaminants has also prompted interest in Pannonibacter species as potential candidates for bioremediation of heavy metal-contaminated environments [1,2]. *P. carbonis* was isolated from coal mine water, whereas *P. anstelovis* was more recently described from bloodstream infections, expanding the known ecological and clinical spectrum of the genus [1–11].

Phenotypically, *Pannonibacter* species are generally aerobic or facultatively anaerobic, oxidase– and catalase-positive, non-fermentative, with species-level differences in substrate utilization and tolerance to temperature, pH, and salinity. From a taxonomic perspective, species delineation within the genus may be challenging because closely related taxa can exhibit high 16S rRNA gene sequence similarity despite substantial genomic divergence. Consequently, whole-genome approaches have become important for resolving species boundaries and identifying previously unrecognized diversity within *Pannonibacter* [1,7–11].

*Pannonibacter* species have been rarely associated with human infections and the few case reports available are mostly in immunocompromised individuals. To date, only eight cases have been documented, including prosthetic valve endocarditis, septicemia, liver abscess, and bacteremia linked to permanent hemodialysis catheter [1–8]. The pathogenic mechanisms of *Pannonibacter* remain poorly understood, but its association with severe infections in immunocompromised hosts suggests an ability to exploit weakened immune defenses [1]. Its adaptability to diverse environments, further underscores the challenges in controlling infections caused by this emerging and opportunistic pathogen [1, 5, 6].

This study characterizes a clinical isolate of *Pannonibacter* from a bacteremia case in Brazil. Comprehensive genomic and phenotypic analyses support its classification as a novel species, for which we propose the name *Candidatus* Pannonibacter alzirae.

## MATERIAL AND METHODS

### Culturing and bacterial identification

The bacterial isolate was received at the Biosafety Level 3 (BSL-3) laboratory of the Aggeu Magalhães Institute. It originated from the patient’s blood culture vial and a subculture on CLED agar, which were referred to our facility following an initial misidentification as *Francisella tularensis*. Within the BSL-3 environment, the isolate was recovered from both the original blood culture vial and the CLED subculture using Tryptic Soy Broth and plated onto MacConkey, chocolate, Müller-Hinton, and Luria–Bertani (LB) agar supplemented with 0.3 g/L L-cystine.

Colonies from both the original specimen and the primary diagnostic culture were subjected to 16S rRNA gene PCR amplification using universal primers (forward 5’-AGAGTTTGATCATGGCTCAG-3’ and reverse: 5’-GGTTACCTTGTTACGACTT-3’). Methodological details are provided in the Supplementary Material. Amplicons were sequenced via Sanger methodology. Resulting sequences were assembled into contigs using BioEdit software (Tom Hall) and analyzed by Basic Local Alignment Tool (BLASTn) against the NCBI non-redundant nucleotide database for taxonomic identification.

Additionally, MALDI-TOF analysis was performed. Colonies were inactivated using formic acid according to the manufacturer’s protocol. To confirm complete inactivation, an aliquot of the extract was plated on LB agar and incubated for 72 hours at 37°C. All procedures were conducted within the BSL-3 facility of the Aggeu Magalhães Institute.

Following inactivation, each extract was spotted onto a steel target plate, overlaid with a matrix solution of α-cyano-4-hydroxycinnamic acid (HCCA; 10 mg/mL in 50% acetonitrile and 2.5% trifluoroacetic acid), and air-dried. Spectral acquisition was performed using a MALDI Biotyper Sirius system (Bruker Daltonics, Germany) with MTB Compass software. The obtained spectra were then compared against the MALDI Biotyper® reference library for microorganism identification, following the producers’ scoring system.

### Morphological characterization

The morphology of the isolate was assessed via macroscopic, light microscopic, and electron microscopic methods. Macroscopic examination was performed on 5% sheep blood agar plates, while light microscopy involved Gram staining.

For scanning electron microscopy (SEM), samples were washed in PBS buffer and post-fixed in 1% osmium tetroxide. They were then dehydrated through a graded ethanol series, critically point-dried using liquid CO₂, and sputter-coated with a thin layer of gold. Observations were made with a Jeol JSM-5600 microscope operated at 15 kV.

For transmission electron microscopy (TEM), samples were washed in 0.1 M sodium cacodylate buffer and post-fixed in a solution of 1% osmium tetroxide, 2 mM calcium chloride, and 0.8% potassium ferricyanide in 0.1 M cacodylate buffer. Following a subsequent wash, the samples were counterstained with 2.5% uranyl acetate, dehydrated through a graded acetone series, and embedded in EMbed 812 resin. Ultrathin sections (70 nm) were obtained using a Leica EM UC7 ultramicrotome, stained with 5% uranyl acetate and 1% lead citrate, and observed with a FEI Tecnai G2 Spirit BioTwin microscope operated at 120 kV.

### Physiological characterization

Physiological characterization was performed by microcultivation in sterile 96-well plates containing 180 μL of LB broth inoculated with 20 μL of bacterial suspension (OD₆₀₀ = 0.10) prepared from LB agar cultures. Growth was monitored by optical density after 24, 48, and 72 hours and evaluated at 4, 8, 18, 30, 37, 42.5, and 45°C, pH 3–12, and final NaCl concentrations of 0.5–20% (w/v), as well as under aerobic, microaerophilic (candle jar), and 5% CO₂ atmospheres at 37°C. Additionally, biofilm formation was assessed using the crystal violet staining assay. Methodological details are provided in the Supplementary Material.

### Whole-genome sequencing and bioinformatic analysis

DNA extraction was performed using the Invitrogen™ PureLink™ Genomic DNA Mini Kit. A genomic library was prepared with the Illumina DNA Prep kit and sequenced on a MiSeq platform using a 600-cycle v3 cartridge (Illumina Inc.) using a paired-end strategy. This sequencing run generated approximately 4.76 million paired-end reads, totaling roughly 2.34 Gb of raw data.

The raw sequencing reads of *Pannonibacter* sp. CBAS 1226 were subjected to a quality control assessment using FastQC v.0.11.8 (https://github.com/s-andrews/FastQC). Low-quality bases were trimmed using Trim Galore v.0.6.5 (https://github.com/FelixKrueger/TrimGalore/releases), and the filtered reads were assembled using the Unicycler v0.4.8 tool, integrated within the BV-BRC platform (https://www.bv-brc.org). Based on the final assembled genome size of 5.51 Mb, the average sequencing depth was estimated at approximately 425-fold coverage. Genome annotation was performed using Prokka v.1.14.6 (https://github.com/tseemann/prokka). Additionally, 26 publicly available *Pannonibacter* genomes and two outgroup genomes were retrieved from the NCBI database for comparative genomic analyses. Genome quality was assessed using CheckM2 v1.1.0 (https://github.com/chklovski/CheckM2). Genome statistics, including genome size, GC content, completeness, contamination, and assembly metrics, were obtained and used to support comparative genomic analyses.

Species classification was performed using Ribosomal Multilocus Sequence Typing (rMLST) (https://pubmlst.org/species-id), with 53 genes encoding bacterial ribosomal protein subunits through the PubMLST online platform (https://pubmlst.org/species-id).

Phylogenomic analyses were performed using the Type Strain Genome Server (TYGS) (https://tygs.dsmz.de). The genome of strain CBAS 1226 was compared against the curated database of type strains, and closely related taxa were identified using MASH. Intergenomic distances were calculated using the Genome BLAST Distance Phylogeny (GBDP) approach, and phylogenomic trees were inferred from the resulting distance matrices with branch support estimated by pseudo-bootstrap resampling. Digital DNA–DNA hybridization (dDDH) values were calculated using the Genome-to-Genome Distance Calculator (GGDC 3.0) implemented within TYGS. Pairwise Average Nucleotide Identity (ANI) values were calculated using FastANI v1.34 (https://github.com/ParBLiSS/FastANI), while Average Amino Acid Identity (AAI) values were estimated using EzAAI v1.2.4 (https://github.com/endixk/ezaai).

Additionally, to reconstruct robust phylogenetic relationships, full-length 16S rRNA gene sequences were extracted from all analyzed genomes and subjected to comparative analysis. The 16S rRNA gene sequences were identified with Barrnap v0.9 (https://github.com/tseemann/barrnap), aligned using MAFFT v7.525, and trimmed with trimAl v1.4 (https://github.com/inab/trimal). To evaluate the robustness of the inferred topology, phylogenetic trees were reconstructed using two complementary methods. A Neighbor-Joining (NJ) tree was generated in MEGA11 (Tamura et al., 2021) using the Kimura two-parameter (K2P) substitution model with 1,000 bootstrap replicates, whereas a Maximum Likelihood (ML) tree was inferred using IQ-TREE v2 (https://github.com/iqtree/iqtree2), with the best-fitting nucleotide substitution model selected by ModelFinder according to the Bayesian Information Criterion (BIC) and branch support estimated from 1,000 ultrafast bootstrap replicates. *Agrobacterium tumefaciens* and *Polymorphum gilvum* were included as outgroups, and the resulting trees were visualized using the Interactive Tree of Life (iTOL) platform (https://itol.embl.de/).

The phylogenetic relationships between *Candidatus* P. alzirae CBAS 1226 and the type or representative strains of all published *Pannonibacter* species were evaluated using complementary 16S rRNA gene– and genome-based approaches. The 16S rRNA gene sequences were aligned with MAFFT, trimmed with trimAl, and analyzed using maximum-likelihood and BIONJ methods. Genome-based relationships were assessed using maximum-likelihood analysis of the GTDB-Tk bac120 marker alignment (https://github.com/ecogenomics/gtdbtk) and the GBDP approach implemented in TYGS. Node support was evaluated using resampling methods, and *Polymorphum gilvum* SL003B-26A1ᵀ was used as the outgroup in all analyses.

Pan-genome analysis was performed to identify gene clusters uniquely present in strain CBAS 1226 relative to its closest phylogenetic relatives. Annotated GFF3 files generated with Prokka for strain CBAS 1226, *P. indicus* DSM 23407, and *P. phragmitetus* DSM 14782 were used as input for Panaroo v1.7.0 (https://github.com/gtonkinhill/panaroo), which was run in strict cleaning mode to minimize the effects of annotation inconsistencies and fragmented genes. Gene presence/absence matrices were generated and used to identify CBAS 1226-specific gene clusters, defined as clusters present in CBAS 1226 and absent from both reference genomes. The corresponding protein sequences were retrieved from the predicted proteome using custom Python scripts based on Biopython SeqIO and subjected to functional annotation with InterProScan v5.78-109.0 (https://github.com/ebi-pf-team/interproscan). InterProScan annotations were integrated with Prokka annotations and the Panaroo presence/absence matrix to generate a comprehensive annotation table including predicted gene names, locus tags, protein identity domains, InterPro entries, Gene Ontology terms, pathway annotations, and the member databases supporting each annotation.

Whole-genome structural comparisons were additionally performed using Proksee (https://proksee.ca). The annotated genome of strain CBAS 1226 was used as the reference, while *P. indicus* DSM 23407 and *P. phragmitetus* DSM 14782 were included for comparative analysis. Circular genome maps were generated using BLAST-based alignments to assess genome synteny, identify conserved and strain-specific genomic regions, and visualize the distribution of genomic features, including coding sequences (CDSs), tRNAs, rRNAs, and CRISPR loci.

The Virulence Factor Database (VFDB) was used to identify potential virulence genes associated with pathogenicity (https://www.mgc.ac.cn/VFs/main.htm). Since no specific virulence database exists for this genus, the genomes were examined for virulence ortholog genes from other bacterial species.

The identification of antibiotic resistance-related features was conducted using the Comprehensive Antibiotic Resistance Database (CARD) (https://card.mcmaster.ca/), ResFinder (https://genepi.food.dtu.dk/resfinder), and InterProScan (https://www.ebi.ac.uk/interpro/search/sequence/). The presence of plasmids in the assembled genome and closely related genomes was evaluated using MOB-suite v.3.0.3 (https://github.com/phac-nml/mob-suite).

### Ethical aspects

This study was approved by the Research Ethics Committee at Aggeu Magalhães Institute (#85921425.2.0000.5190). Since the patient lives in a remote, hard-to-reach community, clinical data was accessed with a waiver of consent by the local Ethics Committee. All patient-related data was anonymized.

## RESULTS

### Case report

An 11-year-old male patient living in a riverside community in Cametá, Pará-Brazil, was admitted to a pediatric oncology hospital with a two-month history of illness. He presented with fever, anasarca, tachycardia, tachypnea, hepatosplenomegaly, anemia, muscle weakness, and sepsis, requiring immediate intervention. Initial investigations at the referring hospital indicated a diagnosis of acute lymphoblastic leukemia. Further epidemiological investigation revealed that the patient had a wild animal as a pet (*Cuniculus paca*), which was reported to carry ticks, and the patient reported tick bites. Additionally, the family frequently consumed wild animal meat.

Upon admission to the children’s oncology hospital on June 3, 2024, the patient was tested for zika, Chagas disease, leptospirosis, visceral leishmaniasis and toxoplasmosis (all negative). Supportive care included packed red blood cell transfusion and oxygen therapy. Empirical antimicrobial therapy was initiated with meropenem and subsequently adjusted based on clinical evolution. The patient received meropenem from admission (Day 1), vancomycin (added on Day 3), and later ciprofloxacin and fluconazole (both added on Day 10). The patient was successfully treated for the infection and was discharged. At the time of the final revision of this manuscript, he remains under outpatient follow-up for ongoing oncological care.

At the local laboratory, the initial blood culture identified *Francisella tularensis* using the Vitek® 2 system (96% compatibility). Subsequently, the same blood culture was streaked onto blood agar and retested with the Vitek® 2 system, this time showing 91% compatibility with *Sphingomonas paucimobilis*. Given the possibility of a highly pathogenic bacterium, the original blood culture and the strain cultured on CLED agar were sent to the BSL-3 at the Aggeu Magalhães Institute (**Supplementary Figure 1**).

### Species Identification of the CBAS 1226

To resolve the initial diagnostic discrepancy between *F. tularensis* and *S. paucimobilis*, the CBAS 1226 was subjected to a comprehensive identification workflow. Colonies obtained from both the original blood culture and the primary isolate recovered at the diagnostic laboratory underwent PCR amplification of 16S rRNA gene, followed by Sanger sequencing. The resulting 1,465-bp sequence showed 100% identity with *Pannonibacter phragmitetus,* contradicting the initial suspicion of *F. tularensis*.

However, detailed inspection of the MALDI-TOF MS spectra revealed inconsistent identification scores among the *P. phragmitetus* reference spectra available in the commercial database. Strain CBAS 1226 yielded highly confident species-level identification scores (>2.0) when compared with strains LMG5414 and LMG5430, whereas only low-confidence (genus-level; score <2.0) or unreliable identification scores were obtained against strains DSM14780, DSM14781, and the type strain *P. phragmitetus* DSM14782 **(Figure 1A–C)**. These inconsistencies suggested that CBAS 1226 might represent a taxon distinct from *P. phragmitetus*, prompting comprehensive genomic, phylogenetic, morphological, and biochemical characterization.

**Figure 1.**
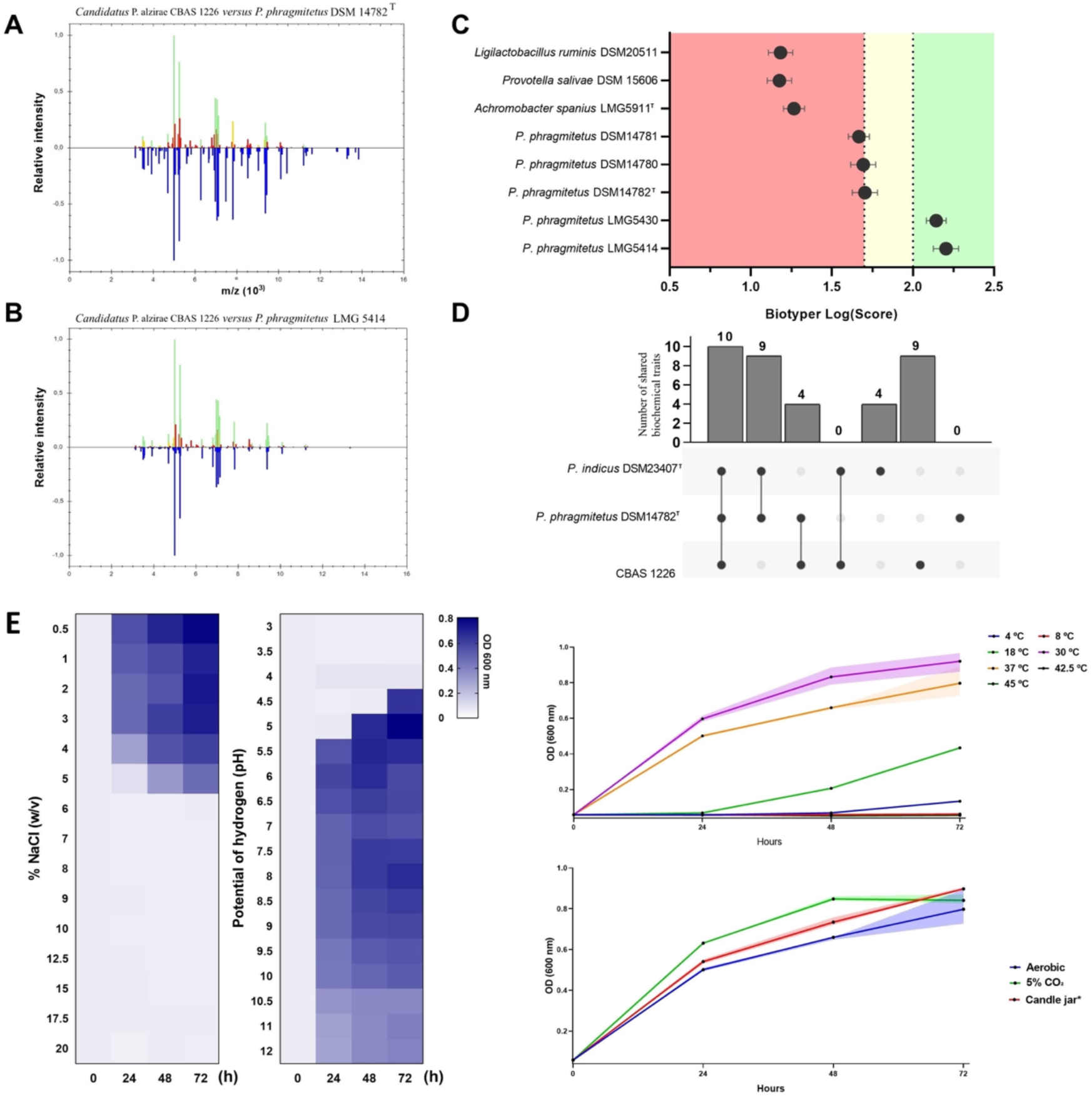
Comparative MALDI-TOF MS and biochemical analysis of *Candidatus* Pannonibacter alzirae CBAS 1226. Legend: (**A** and **B**) MALDI-TOF MS spectrum of *Candidatus* P. alzirae CBAS 1226. In the spectra, the blue color in the lower section represents stored reference spectra from the Compass software library used for matching. The green peaks in the upper section indicate excellent matches, while red and yellow peaks denote mismatched and intermediate matches, respectively. The MALDI-TOF MS analysis identified *P. phragmitetus* with a log score >2.00, with the most intense signals observed at 4,250 and 8,000 Da. (**C**) Comparison of protein spectra of *Candidatus* P. alzirae with the main *Pannonibacter* species available in the MALDI Biotyper database. (**D**) Biochemical comparison of *Candidatus* P. alzirae CBAS 1226 with *P. phragmitetus* DSM 14782T and *P. indicus* DSM 23407, highlighting shared and distinct metabolic traits. (**E**) Physiological characterization of *Candidatus* P. alzirae CBAS 1226 under different pH, NaCl, temperature, and atmospheric conditions.

### Genomic analysis

The whole-genome sequencing produced a high-quality assembly with 38 contigs and a N50 value of 530,907 bp. CheckM2 quality control assessments confirmed approximately 100% of completeness. **Supplementary Table S1** summarizes the genomic characteristics of the other strains used in this study.

Whole-genome phylogenomic analysis consistently placed strain CBAS 1226 within a distinct and well-supported lineage, clearly separated from the type strains of *P. phragmitetus*, *P. indicus*, and the remaining recognized members of the genus **(Figure 2A)**. Likewise, phylogenetic reconstruction based on the 16S rRNA gene using the TYGS, Neighbor-Joining (NJ), and Maximum Likelihood (ML) approaches yielded congruent topologies (**Supplementary Figures 2 and 3**). Although minor differences were observed in the branching order of closely related reference strains, all three methods consistently recovered strain CBAS 1226 as a distinct lineage within the genus (**Figure 2B**). rMLST analysis further supported this distinction by identifying *P. phragmitetus* and *P. indicus* as the closest taxa, although with low assignment support (26% and 18%, respectively), indicating that CBAS 1226 could not be confidently assigned to any currently recognized species (**Supplementary Table S2**). Further phylogenomic evidence for this distinction is presented in **Supplementary Figure 4**.

**Figure 2.**
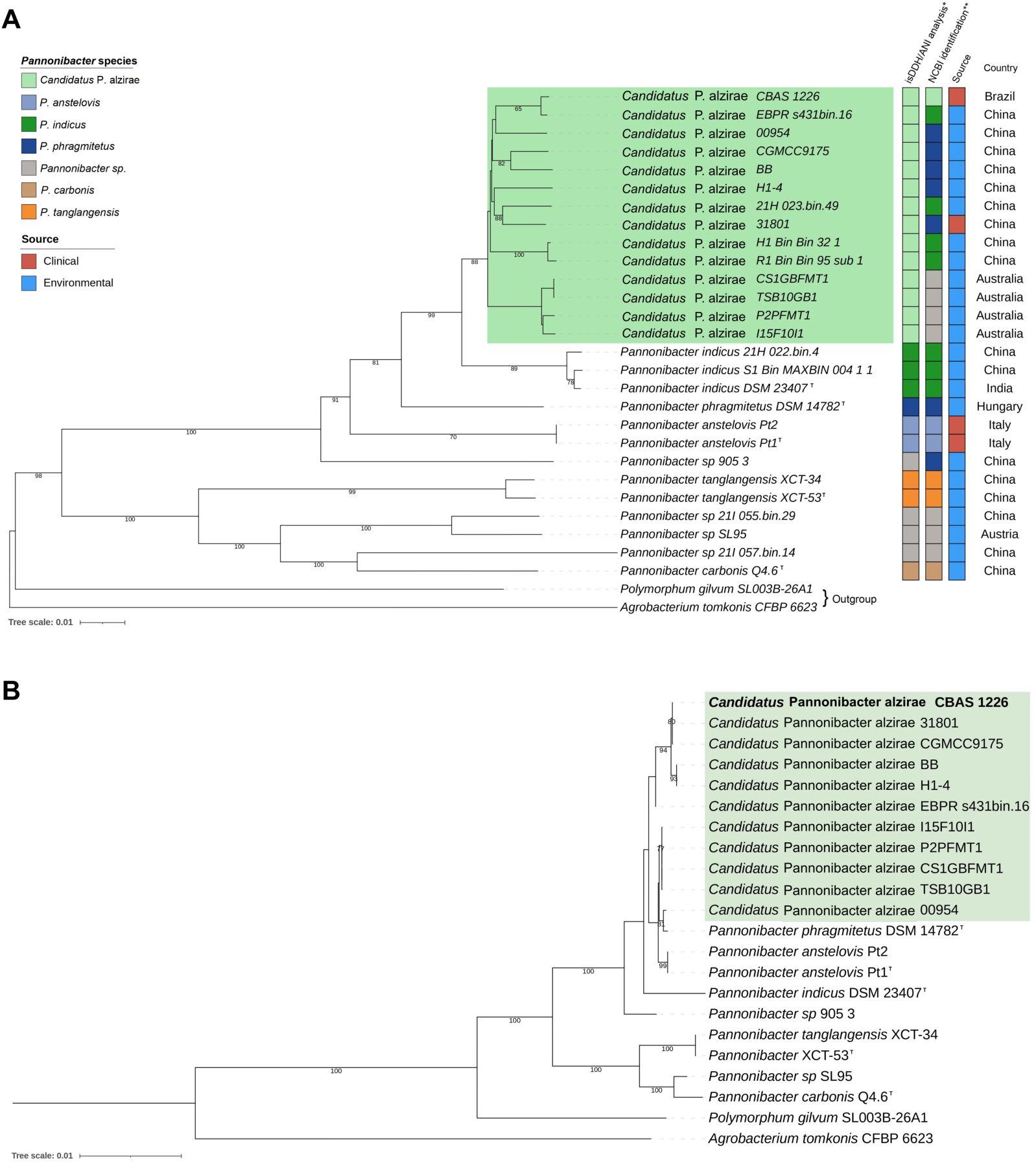
Phylogenetic relationships among *Pannonibacter* species. Legend: **(A)** Whole-genome phylogenomic tree inferred using the Genome BLAST Distance Phylogeny (GBDP) approach implemented in the Type Strain Genome Server (TYGS). *Candidatus* Pannonibacter alzirae strain CBAS 1226 forms a distinct and well-supported lineage, clearly separated from the type strains of *P. phragmitetus*, *P. indicus*, and the remaining recognized *Pannonibacter* species included in the analysis. **(B)** Phylogenetic tree based on 16S rRNA gene sequences. The tree was inferred from GBDP distances using the CharacterCoverage algorithm and distance formula d5, followed by balanced minimum-evolution analysis with FastME v2.1.6.1. Branch support was estimated from 100 pseudo-bootstrap replicates. *Candidatus* P. alzirae CBAS 1226 is highlighted in bold, and the *Candidatus* P. alzirae cluster is shaded in green. Some strains could not be included because complete or sufficiently high-quality 16S rRNA gene sequences were not recovered from their genome assemblies, a common limitation of metagenome-assembled genomes (MAGs), in which ribosomal operons may be fragmented or absent due to assembly and binning processes. Nevertheless, these genomes were included in the whole-genome phylogenomic analysis, which provided higher taxonomic resolution than the single-locus 16S rRNA gene analysis. In both analyses, the *Candidatus* P. alzirae CBAS 1226 lineage was consistently recovered as a distinct and well-supported clade.

**Figure 3.**
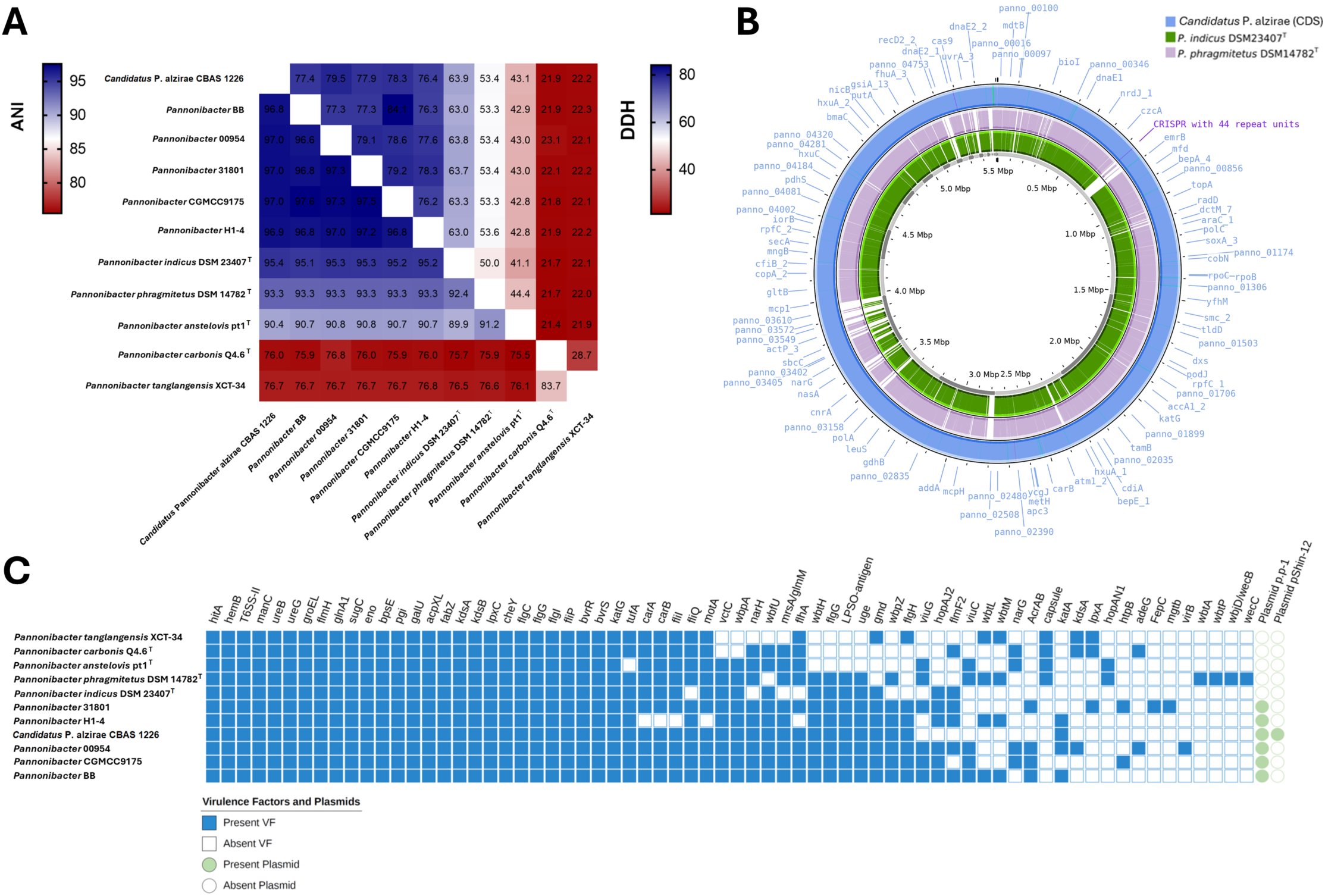
Genomic relatedness of *Candidatus* Pannonibacter alzirae CBAS 1226 to closely related species. Legend: (**A**) dDDH and ANI values were below the respective species delineation thresholds when strain CBAS 1226 was compared with the closest type strains, supporting its classification as a novel species within the genus *Pannonibacter*. Additional information can be found at **Supplementary Table S6**. (**B**) The outermost ring (blue) represents the coding sequences (CDS) of *Candidatus* P. alzirae CBAS 1226 displayed in circular coordinates. Genomic positions are indicated by the scale (Mbp) along the inner circumference. Selected annotated genes are labeled outside the circle; a CRISPR array containing 44 repeat units is highlighted in magenta. The middle ring (lavender) and the innermost ring (green) display syntenic regions between *Candidatus* P. alzirae CBAS 1226 and the reference strains *P. phragmitetus* DSM14782T and P. indicus DSM23407T, respectively. Colored arcs indicate genomic regions of the reference strains with synteny to *Candidatus* P. alzirae CBAS 1226, whereas gaps represent regions of the CBAS 1226 genome absent or highly divergent in the reference strains. The figure was generated using Proksee (https://proksee.ca). **C)** Functional annotation identified multiple antibiotic resistance genes and genome overview showing the two plasmids absent in other *Pannonibacter* species.

Comparative genomic analyses summarized in the ANI/dDDH matrix (**Figure 3A**) revealed dDDH values ranging from 43.1% to 63.9% between strain CBAS 1226 and the type strains of its closest relatives, remaining below the 70% threshold for species delineation. Correspondingly, ANI values ranged from 90.4% to 95.4%. The highest ANI was observed with *P. indicus* DSM 23407 (95.4%), placing this comparison at the lower boundary of the accepted species cutoff (95–96%), whereas values for *P. phragmitetus* DSM 14782 (93.3%) and *P. anstelovis* Pt1 (90.4%) were clearly below this threshold. In contrast, AAI values remained consistently high (∼97%) among the closest *Pannonibacter* species (**Supplementary Table S6**), indicating strong conservation of shared orthologous proteins while remaining consistent with the phylogenomic and whole-genome similarity analyses.

Whole-genome comparison between strain CBAS 1226 and the genomes of its closest type strains, *P. indicus* DSM 23407 and *P. phragmitetus* DSM 14782, revealed extensive conservation of genome organization (**Figure 3B**). Despite the overall conservation of gene order, several genomic regions were unique to strain CBAS 1226. Pan-genome analysis identified 537 gene clusters exclusive to CBAS 1226, of which approximately half encoded hypothetical proteins. The annotated fraction was enriched in genes associated with mobile genetic elements and defense systems, including Type IV secretion, CRISPR-Cas, restriction-modification, and toxin-antitoxin modules. Additional strain-specific loci comprised genes related to heavy-metal homeostasis, heme and iron acquisition, multidrug efflux transport, and a complete NiFe hydrogenase gene cluster, further highlighting the distinct accessory genome of strain CBAS 1226 (**Supplementary Table S7**).

Collectively, these findings consistently distinguish strain CBAS 1226 from its closest phylogenetic relatives and demonstrate that it represents a genomically distinct lineage within the genus *Pannonibacter*. The convergence of phylogenetic, comparative genomic, and pan-genome analyses, together with the phenotypic characteristics described herein, provides robust support for the recognition of the novel species *Candidatus* Pannonibacter alzirae.

### Morphological aspects

After 48 hours of incubation on blood agar at 30°C, *Candidatus* P. alzirae formed smooth, slightly convex colonies, measuring 0.5 to 1 mm in diameter. The colonies displayed an opaque grayish-white coloration with well-defined edges, with no evidence of pigmentation or hemolytic activity (**Figure 4A**). Optical and electron micrographs revealed Gram-stain-negative rod-shaped cells measuring between 2 and 4 µm in length and 0.3 to 0.6 µm in width, with irregular surfaces. Some cells exhibited a lateral flagellar structure (**Figure 4 B-D**).

**Figure 4.**
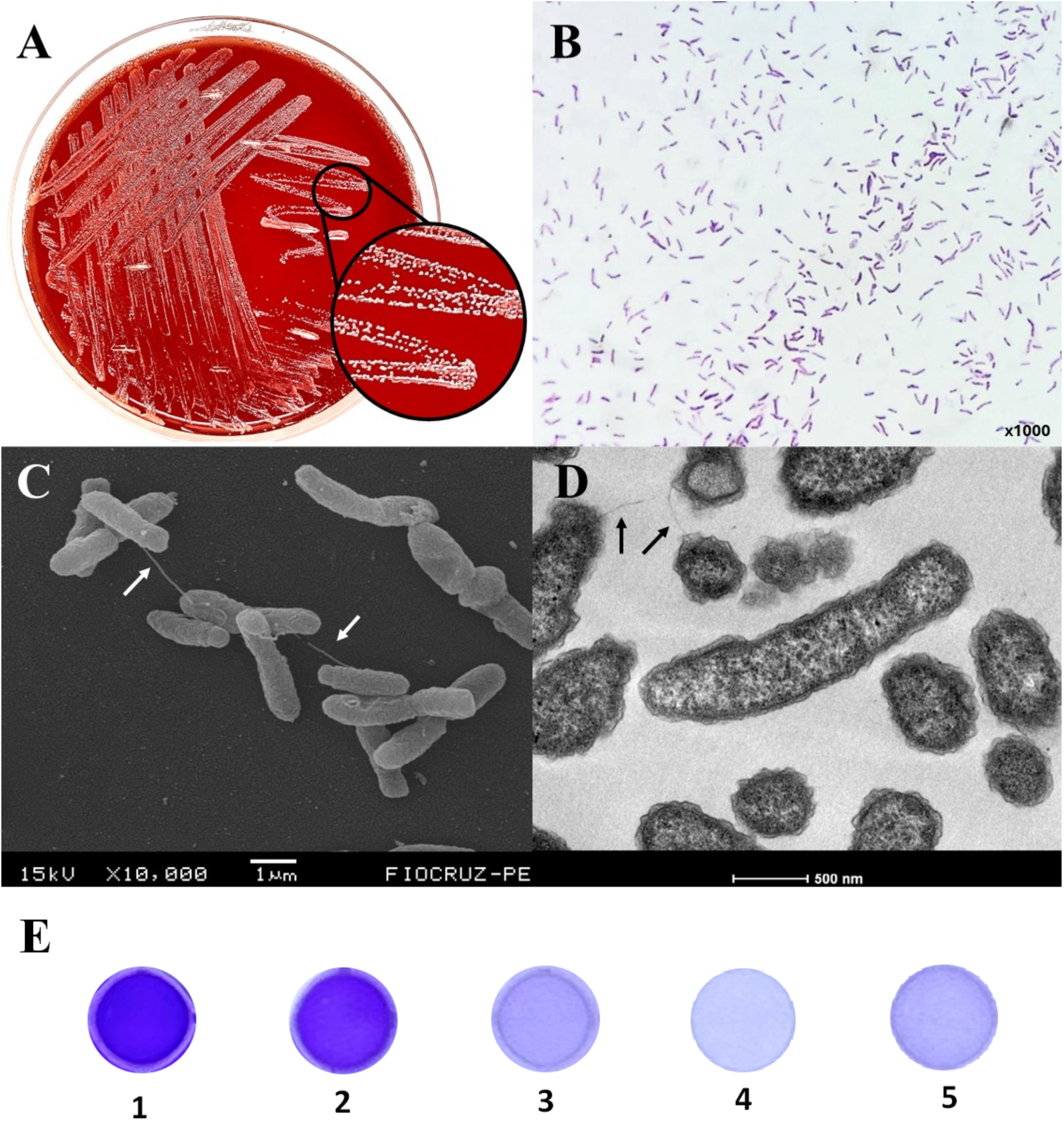
Colony morphology and microscopic characterization of *Candidatus* Pannonibacter alzirae CBAS 1226. Legend: (**A**) Opaque grayish-white colonies with well-defined edges and a shiny surface texture on blood agar medium. (**B**) Optical microscopy (Gram staining) showing rod-shaped Gram-stain-negative cells. (**C**) Scanning electron microscopy (SEM) image highlighting rod-shaped cells, with some exhibiting a flagellar structure at one end (arrow). (**D**) Transmission electron microscopy (TEM) image showing a flagellar structure at one cell end (arrow). (**E**) Crystal violet biofilm assay illustrating the reference categories of biofilm formation. 1 = positive control (strong); 2 = positive control (moderate); 3 = positive control (weak); 4 = negative control (non-biofilm producer); 5 = *Candidatus* P. alzirae CBAS 1226, classified as a weak biofilm producer.

### Bacterial traits, antimicrobial resistance and biofilm formation

*Candidatus* P. alzirae was characterized for 82 distinct biochemical tests (**Supplementary Table S3**) and exhibited 21 shared test results with *P. phragmitetus* DSM 14782 and 9 with *P. indicus* DSM 23407. Using biochemical assays, *Candidatus* P. alzirae CBAS 1226 can be distinguished from the *P. indicus* and *P. phragmitetus* type by nine biochemical traits: alkaline phosphatase, β-glucosidase, D-cellobiose as carbon source (CEL), D-fructose as carbon source (FRU), D-glucose as carbon source (GLU), D-mannose as carbon source (MNE), D-melibiose as carbon source (MEL), esculin as carbon source (ESC) and L-arginine (**Figure 1D**).

Overall, carbapenems and fluoroquinolones exhibited low MICs. Moderate MIC values were observed for ceftazidime and cefepime and higher MICs were noted for piperacillin-tazobactam, amoxicillin-clavulanic acid and aztreonam (**Supplementary Table S4**). *Candidatus* P. alzirae CBAS 1226 exhibited weak biofilm-forming capacity under the conditions tested (mean OD₅₇₀ = 0.103 ± 0.024; **Figure 4E**).

### Physiological aspects

*Candidatus* P. alzirae grew over a broad range of environmental conditions (**Figure 1E**). Growth was detected at 18, 30, and 37°C after 24 h. No growth was detected at 42.5, or 45°C. *Candidatus* P. alzirae tolerated NaCl concentrations up to 5% (w/v), with no growth observed at higher concentrations. Growth occurred under aerobic, microaerophilic (candle jar), and 5% CO₂ atmospheres. Growth was observed from pH 4.5 and throughout the alkaline pH range tested (pH 12), whereas no growth was detected at pH ≤ 4.0.

### Genomic aspect of *Candidatus* Pannonibacter alzirae

*Candidatus* P. alzirae CBAS 1226 had a genome size of 5.5 Mb (whereas *P. indicus* and *P. phragmitetus* type strains had 4.2 and 4.8, respectively), and a GC content of 63.4% (**Table 1**), and two plasmids: p.p-1 (268,093 bp, non-mobilizable) and pShin-12 (48,304 bp, conjugative), that are not found in other species within the genus (**Figure 3C**). InterProScan identified beta-lactamases, multidrug efflux pumps, aminoglycoside phosphotransferases, and tetracycline resistance proteins (**Supplementary Table S5**). Sixty-eight genes potentially involved in virulence were identified, encompassing those related to toxin secretion (*hitA*), production of exopolysaccharides (*bpsE*), iron acquisition (*hemB*), stress resistance (*katG*, *groEL*), lipopolysaccharide biosynthesis (*lpxC*, *acpXL*, *wbpA*) and biofilm formation (*flgC*, *flgG*, *fliP*, *motA*) (**Figure 3C**).

**Table 1:** Phenotypic and genomic characteristics differentiating *Candidatus* Pannonibacter alzirae from its closest related type strains.

| **Characteristic** | ***Candidatus* P. Alzirae CBAS 1226** | ***P. phragmitetus* DSM 14782<sup>T</sup>** | ***P. indicus* DSM 23407<sup>T</sup>** |
|---|---|---|---|
| **Origin of strain** | Human (bacteremia) | Environmental (decomposing reed rhizomes) | Environmental (hot spring sediment) |
| **Colony (blood agar, 48h, 30 °C)** | Greyish-white, non-pigmented, non-hemolytic | Data not provided | Data not provided |
| **Optimal growth temperature** | 30 to 37°C | 22 to 28 °C | 30 °C |
| **Maximum [NaCl] tolerated** | 5% | 5% | ND |
| **pH range** | 4.5 to ≥ 12 | 7 to 11 | 6 to 11 |
| **Enzyme production** |  |  |  |
| Alkaline phosphatase | - | + | + |
| α-Glucosidase | + | + | - |
| β-Glucosidase | + | - | - |
| L-Arginine | - | + | + |
| L-Rhamnose | - | + | ND |
| Lysine decarboxylase | - | ND | + |
| **Carbon source for growth** |  |  |  |
| 2-ketoglutarate | + | - | ND |
| 5-ketogluconate | - | + | ND |
| D-Arabitol | - | - | + |
| D-Cellobiose | - | + | + |
| D-Fructose | - | + | + |
| D-Galactose | - | + | ND |
| D-Galacturonate | - | + | ND |
| D-Glucose | - | + | + |
| D-Mannose | - | + | + |
| D-Trehalose | - | + | ND |
| Esculin | - | + | + |
| L-Arabinose | - | + | ND |
| L-Proline | + | - | ND |
| L-Rhamnose | - | + | ND |
| Maltose | - | + | ND |
| Palatinose | - | + | ND |
| Sucrose | - | + | ND |
| **Acid production** |  |  |  |
| D-Glucose | - | - | + |
| **DNA G+C content (%)** | 63.4 | 63 | 63.5 |
| **Genome size (Mb)** | ~5.5 | ~4.8 | ~4.2 |
| **Plasmid content** | p.p-1 (268 kb) and pShin-12 (48 kb) | ND | ND |

Two extrachromosomal structures were found: a large plasmid-like contig of 268 kb (recovered across seven contigs) showing similarity to previously described *Pannonibacter* plasmids, and a 48 kb conjugative plasmid (recovered as a single contig) carrying MOBQ relaxase and MPF_T signatures consistent with the *Rhizobiaceae* family.

### Nomenclatural status of the proposed taxon

The novel taxon described in this study is proposed under the *Candidatus* designation because it does not currently fulfil the requirements for valid publication of a species name under the ICNP. Owing to current Brazilian national legal and regulatory restrictions governing the international deposition of biological materials, the proposed type strain cannot presently be deposited in two publicly accessible culture collections located in different countries, as required by the ICNP. Consequently, the proposed name does not currently have standing in prokaryotic nomenclature. The *Candidatus* designation therefore reflects a conjectural rather than a taxonomic limitation.

## DISCUSSION

Reported cases of infection caused by *Pannonibacter* are exceedingly rare, mostly documented in immunocompromised individuals [2, 7], which aligns with the current report in an acute lymphoblastic leukemia patient. The initial misidentification of *Pannonibacter* as *F. tularensis* using biochemical/phenotypical or low-resolution molecular methods highlights a critical limitation of these tools in cases involving rare Gram-stain-negative bacteria [10]. Similar challenges have been documented in other cases of *Pannonibacter* sp.,where definitive identification was only achieved through high resolution molecular and genomic tools [1].

The dDDH and ANI values, which were below the respective species delineation thresholds, together with the distinct biochemical and proteomic features, indicate that *Candidatus* P. alzirae represents a novel taxon within the genus. These findings are consistent with the growing recognition of genomic diversity among *Pannonibacter* species [1, 7]. Notably, *Candidatus* P. alzirae clusters with other strains currently assigned as *P. phragmitetus* in deposit information, forming a group that exhibits significant genetic divergence from the *P. phragmitetus* type strain. The consistent clustering of these strains in phylogenetic analyses underscores the need for a comprehensive reevaluation of the genus using polyphasic taxonomic approaches.

The genus *Pannonibacter* remains relatively underrepresented in public genome databases despite its increasing environmental and clinical relevance. Recent taxonomic studies have demonstrated that species boundaries within the genus cannot always be reliably resolved using the 16S rRNA gene alone. This limitation became evident during the description of *Pannonibacter anstelovis*, in which isolates initially assigned to *P. phragmitetus* based on 16S rRNA sequence similarity were subsequently shown, through whole-genome comparisons (ANI, dDDH, and phylogenomics), to represent a distinct species [1]. Likewise, retrospective examination of strain 31801, originally described as *P. phragmitetus*, reveals a similar pattern, despite its high similarity to strain CGMCC9175, its ANI relative to the authentic type strain *P. phragmitetus* DSM 14782 already fell below the currently accepted species threshold [11].

Synteny analysis showed that more than 537 genes were exclusively found in *Candidatus* P. alzirae CBAS 1226, including those encoding the T4SS, toxin-antitoxin systems, and CRISPR-Cas proteins. These genes form clusters in the genome, supporting the hypothesis that their acquisition was from a single accessory genomic island or conjugative element through horizontal gene transfer.

The physiological characterization of the novel species revealed broad growth versatility, with the ability to grow at temperatures ranging from 18 to 37 °C, under highly alkaline conditions, and across different oxygen availability conditions. This finding is in line to the reports from other species from the same genus isolated in alkaline, cold-temperature lakes [3]. It is important to note that the biochemical characterization from CBAS 1226 strain was performed using semi-automated diagnostic assays, while data from other species were obtained from public databases, which represents a methodological limitation and should be interpreted with caution.

The antimicrobial susceptibility testing of *Candidatus* P. alzirae CBAS 1226 was consistent with previous reports of *Pannonibacter* species, and the resistance determinants identified by WGS provided a genetic basis for this phenotypic resistance [1, 11, 12]. In the present case, the empirical antimicrobial regimen was consistent with the isolate’s in vitro susceptibility. Phenotypic testing confirmed high susceptibility to meropenem and ciprofloxacin, and whole-genome analysis corroborated the absence of acquired resistance genes to these drug classes. The detected intrinsic beta-lactamase-like domains and efflux systems align with the observed resistance to penicillins and aztreonam, underscoring a genotype-phenotype concordance that validated the therapeutic approach [1, 11, 12].

The detection of conjugative plasmids and resistance genes raises concerns about the potential for emerging resistance under selective pressure [13–15]. Although long-read sequencing would be necessary to resolve the complete structure and circularize the larger plasmid, the 425× coverage achieved with short-read technology was adequate for plasmid detection and characterization within the taxonomic scope of this study. The identification of genes related to the Type VI secretion system (T6SS-II) and urease production (*ureB*, *ureG*) underscores its potential to cause invasive disease, particularly in immunocompromised hosts [11, 16].

## CONCLUSION

This study describes *Candidatus* Pannonibacter alzirae, a novel *Pannonibacter* species isolated from an immunocompromised pediatric patient in Brazil. A polyphasic taxonomic approach combining phenotypic characterization with comparative genomic analyses demonstra ted that strain CBAS 1226 represents a distinct evolutionary lineage within the genus. In addition to supporting the description of a novel species, our results indicate that several publicly available genomes currently assigned to *P. phragmitetus* likely belong to *Candidatus* Pannonibacter alzirae, emphasizing the importance of genome-based taxonomic reassessment. The presence of plasmids, virulence-associated genes, and the ability to grow under diverse environmental conditions further expand current knowledge of the biology and opportunistic potential of this species. Together, these findings broaden the known diversity of the genus *Pannonibacter* and highlight the value of integrating genomic and phenotypic approaches for the accurate characterization of uncommon clinical isolates.

## DESCRIPTION OF *Candidatus* Pannonibacter alzirae sp. nov

*Candidatus* Pannonibacter *alzirae* (al.zi’rae. N.L. gen. fem. n. alzirae, of Alzira, named in honour of Dr. Alzira Maria Paiva de Almeida, a pioneering scientist in the Department of Microbiology of the Aggeu Magalhães Institute, Brazil).

Cells are Gram-stain-negative, non-spore-forming rods, approximately 2–4 µm long and 0.3–0.6 µm wide, and motile by means of a lateral flagellum. Colonies on blood agar are smooth, slightly convex, opaque, greyish-white, non-pigmented, non-haemolytic and 0.5–1 mm in diameter after incubation at 30 °C for 48 h. Growth occurs under aerobic, microaerophilic and 5 % CO₂ atmospheres and at 18 to 37 °C, pH 4.5 to ≥ 12, and in the presence of 5% NaCl (w/v), with optimal growth at 30–37 °C, pH 5.0 –9.0 and NaCl concentration of 0.5 to 4%.

Produces catalase, oxidase, α-glucosidase and β-glucosidase, but not alkaline phosphatase or lysine decarboxylase. Uses L-proline and 2-ketoglutarate as carbon sources, but not D-arabitol, D-cellobiose, D-fructose, D-galactose, D-galacturonate, D-glucose, D-mannose, D-trehalose, L-arabinose, L-arginine, L-rhamnose, 5-ketogluconate, maltose, palatinose or sucrose. Does not hydrolyse esculin. Does not produce acid from D-glucose. Comparative biochemical and genomic characteristics are summarized in **Table 1 and Supplementary Table S8**.

The CBAS 1226 strain (reference) was isolated from the blood of an 11-year-old male patient with acute lymphoblastic leukemia and bacteremia in Cametá, Pará, Brazil. The DNA G+C content of the genome of the reference strain is 63.4 mol%, with an approximate genome size of 5.5 Mb. The Whole Genome Shotgun project for strain CBAS 1226 has been deposited in DDBJ/ENA/GenBank under accession number **JBVOUO000000000** (version JBVOUO010000000; BioProject PRJNA1223427, BioSample SAMN46819472, SRA SRR32335018). The 16S sequence coordinate within the Whole-genome sequence is **JBVOUO010000026.1**. The live strain was deposited in the Bacteria Collection from Environment and Health (CBAS), FIOCRUZ, Brazil.

## Supporting information

Supplementary Material for "Candidatus Pannonibacter alzirae sp. nov. isolated from a septic immunocompromised patient in Brazil"

## ACKNOWLEDGEMENTS

We thank the Núcleo de Plataformas Tecnológicas (NPT) at the Aggeu Magalhães Institute for their support, as well as the Multiservice Laboratory for Research and Diagnosis in Tropical Diseases (LPDDT) for mass spectrometry analyses. We also extend our gratitude to the Brazilian Ministry of Health for institutional support; the Pará State Center for Strategic Health Surveillance Information (CIEVS-PA) and the Pará State Public Health Reference Laboratory (LACEN-PA) for epidemiological and diagnostic collaboration; and the Octávio Lobo Children’s Oncology Hospital (Belém, Pará) for patient care and sample referral.

## DATA AVAILABILITY

The underlying data for this article is available within the manuscript. The strain was deposited in the Fiocruz/CBAS collection (https://cbas.fiocruz.br, identification: CBAS 1226). The Whole Genome Shotgun project for *Candidatus* P. alzirae CBAS 1226 has been deposited at DDBJ/ENA/GenBank under the accession JBVOUO000000000. The version described in this paper is version JBVOUO010000000. The raw sequencing data are available in the NCBI Sequence Read Archive database under SRA accession number SRR32335018, associated with BioProject PRJNA1223427 (BioSample SAMN46819472). The 16S rRNA gene sequence of CBAS 1226 was extracted directly from the complete genome assembly using BLAST-based searches against reference 16S rRNA gene sequences. A comprehensive list of accession numbers for the publicly available genome assemblies used for comparative purposes, as well as for the extracted 16S rRNA gene sequences, is provided in **Supplementary Table S1**. The table also includes the genomic coordinates of the 16S rRNA gene within the complete genome assembly.

### Authors’ contribution

I.V.R. –Conceptualization, Data curation, Formal analysis, Investigation, Methodology, Writing –original draft; B.S.T.M. (Bioinformatics analysis, Methodology, Writing –original draft); J.L.L.P.P. (Bioinformatics analysis); G.L.W. (Bioinformatics analysis); T.L.C. (Bioinformatics analysis, Writing –review & editing); C.A.N.A. (Mass spectrometry analysis); R.G.L.N. (Mass spectrometry analysis); K.L.A.S. (Scanning and transmission electron microscopy analysis); M.E.B.A.C. (Biofilm assay, Physiological characterization); M.S., M.L.N. and A.C.M.F. (Project administration, Writing –review & editing); M.F.B. (Conceptualization, Data curation, Formal analysis, Investigation, Methodology, Writing –original draft).

