## Supplementary Material for "Candidatus Pannonibacter alzirae sp. nov. isolated from a septic immunocompromised patient in Brazil" for "*Candidatus* Pannonibacter alzirae sp. nov. isolated from a septic immunocompromised patient in Brazil"

Rocha & Melo et al., 2026

**SUPPLEMENTARY MATERIAL**

### 1 SUPPLEMENTARY METHODS

#### 1.1 PCR amplification of the 16S rRNA gene

The 16S rRNA gene was amplified by PCR using the universal primers 8F (5′-AGAGTTTGATCATGGCTCAG-3′) and 1492R (5′-GGTTACCTTGTTACGACTT-3′), generating an amplicon of 1,465 bp [1]. PCR reactions were performed in a final volume of 25 µL containing 1× GoTaq® Green Master Mix (Promega), 50 ng of genomic DNA, and 0.4 µM of each primer.

Amplification was carried out with an initial denaturation at 95°C for 5 min, followed by 35 cycles of denaturation at 95°C for 45 s, annealing at 55°C for 45 s, and extension at 72°C for 2 min, with a final extension at 72°C for 5 min. PCR products were analyzed by electrophoresis on a 1.2% agarose gel.

#### 1.2 Assessment of biofilm formation

Biofilm formation by *Candidatus* P. alzirae was evaluated using the crystal violet staining method, adapted from a previously described protocol [2]. An overnight culture grown in LB broth was adjusted to an $OD_{600}$ of 0.08–0.13. Aliquots of 180 µL of LB supplemented with 1% glucose and 20 µL of the bacterial suspension were dispensed, in triplicate, into a sterile 96-well flat-bottom polystyrene microplate. Following incubation at 37°C for 24 h, the wells were washed three times with sterile deionized water and stained with 200 µL of 0.1% crystal violet for 20 min. Excess stain was removed by washing the wells three times with sterile deionized water, and the retained dye was solubilized with 200 µL of 95% ethanol for 20 min at room temperature. Biofilm biomass was quantified by measuring the absorbance at 570 nm using a Benchmark Plus Microplate Reader (Bio-Rad).

The cut-off optical density (ODc) was defined as the mean OD of the negative control (LB medium only) plus three standard deviations (SD). Biofilm-forming capacity was classified as follows: OD ≤ ODc, non-producer; ODc < OD ≤ 2 × ODc, weak producer; 2 × ODc < OD ≤ 4 × ODc, moderate producer; and OD > 4 × ODc, strong producer [3].

#### 2 SUPPLEMENTARY FIGURES

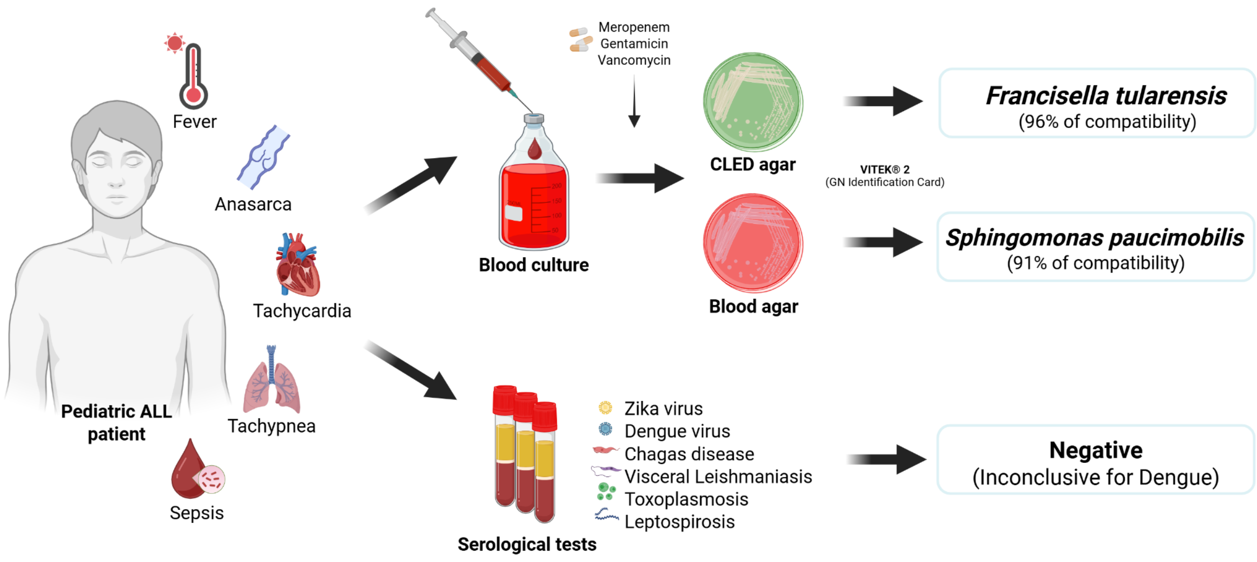


**Supplementary Figure 1: Sample collection overview.**

A

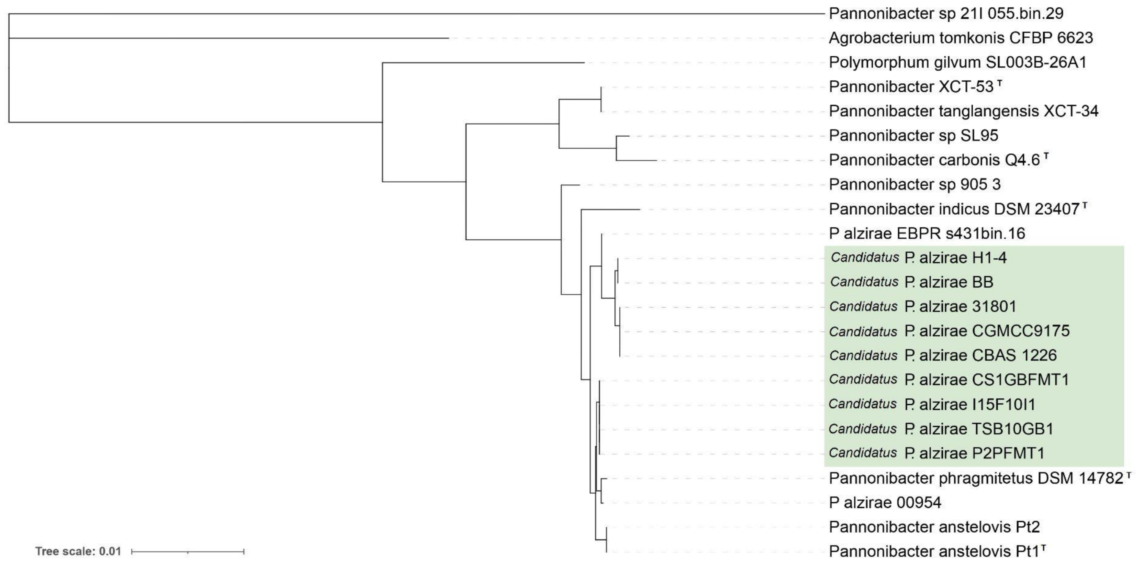


B

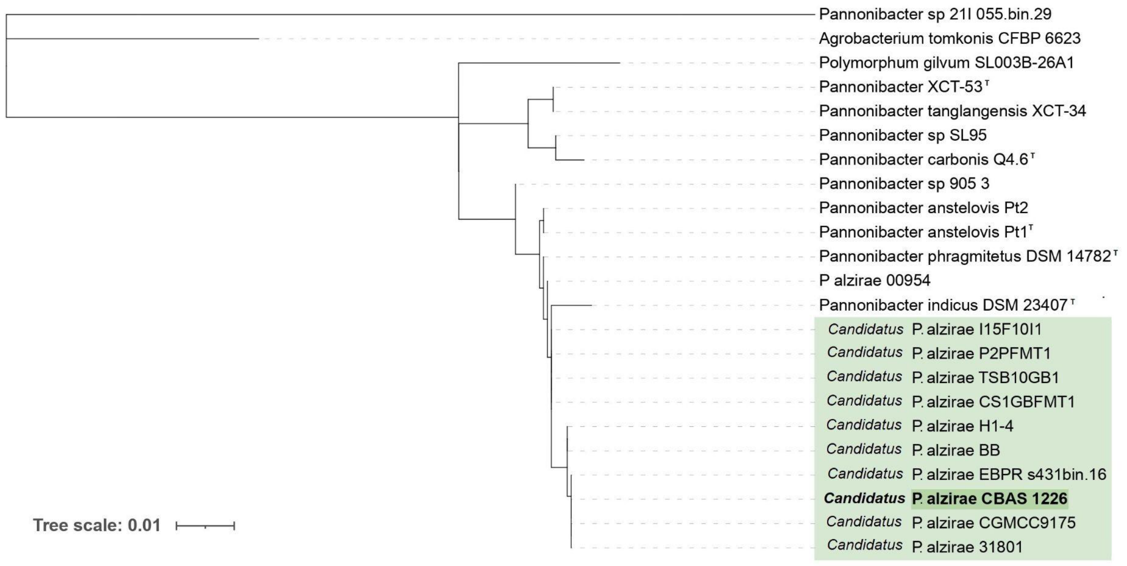


**Supplementary Figure 2: Phylogenetic reconstruction based on 16S rRNA gene sequences of *Pannonibacter* strains included in this study.** (**A**) Neighbor-Joining (NJ) tree reconstructed in MEGA11 using the Kimura two-parameter (K2P) substitution model with 1,000 bootstrap replicates. (**B**) Maximum Likelihood (ML) tree inferred using IQ-TREE v2 with the best-fitting nucleotide substitution model selected by ModelFinder according to the Bayesian Information Criterion (BIC) and branch support estimated from 1,000 ultrafast bootstrap replicates. *Agrobacterium tomkonis* CFBP 6623 and *Polymorphum gilvum* SL003B-26A1 were included as outgroups. Genomes proposed to belong to *Candidatus* Pannonibacter alzirae sp. nov. are highlighted in green.

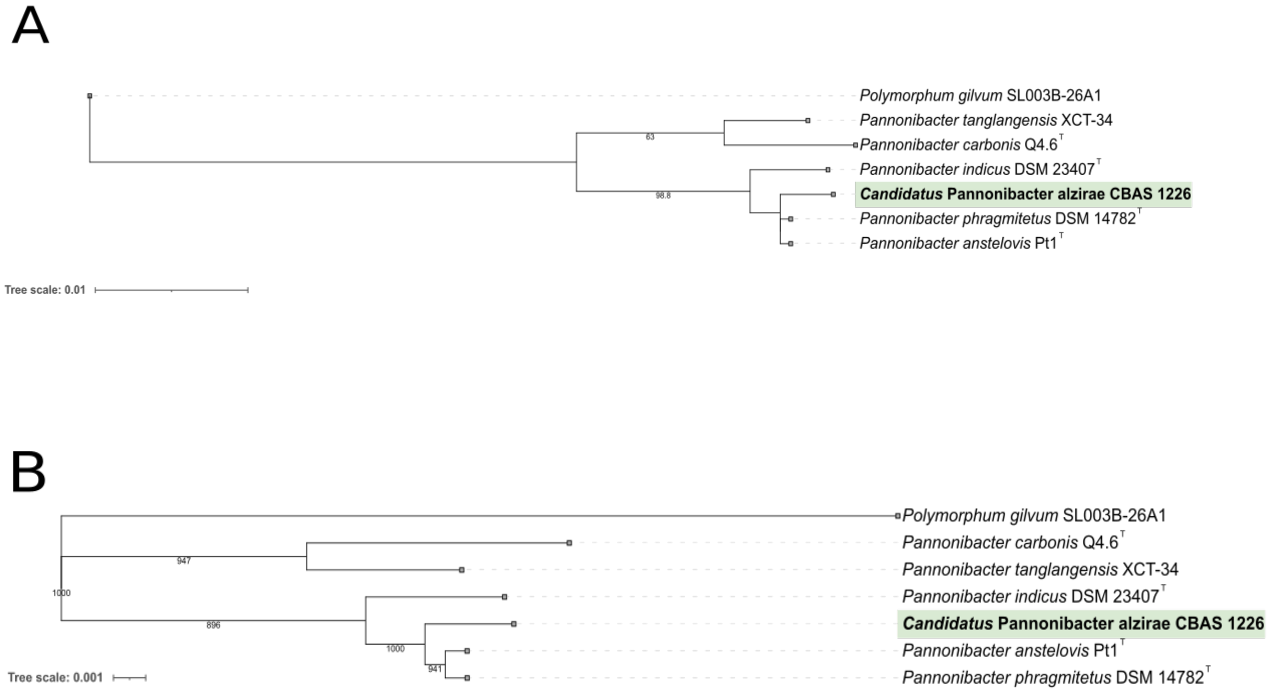


**Supplementary Figure 3: Phylogenetic relationships among Pannonibacter type strains based on the 16S rRNA gene, inferred using two independent methods.** **(A)** Maximum-likelihood (ML) tree inferred with IQ-TREE 2 under the best-fitting nucleotide substitution model (HKY+F+I), selected according to the Bayesian information criterion (BIC). Node support was assessed using the SH-like approximate likelihood ratio test (SH-aLRT) with 1,000 replicates; only support values ≥50% are shown. **(B)** Distance-based tree inferred using the BIONJ algorithm based on Kimura two-parameter distances, as implemented in the R package *ape*. Node support was assessed using 1,000 bootstrap replicates, and support values are shown as percentages at internal nodes. The dataset included the type strains of all published *Pannonibacter* species and CBAS 1226, representing the novel taxon *Candidatus* P. alzirae described in this study. *Polymorphum gilvum* SL003B-26A1$^{T}$ was used as the outgroup in both analyses. Branch lengths represent nucleotide substitutions per site, with scale bars indicated separately for each panel. Both methods recovered the same overall topology.

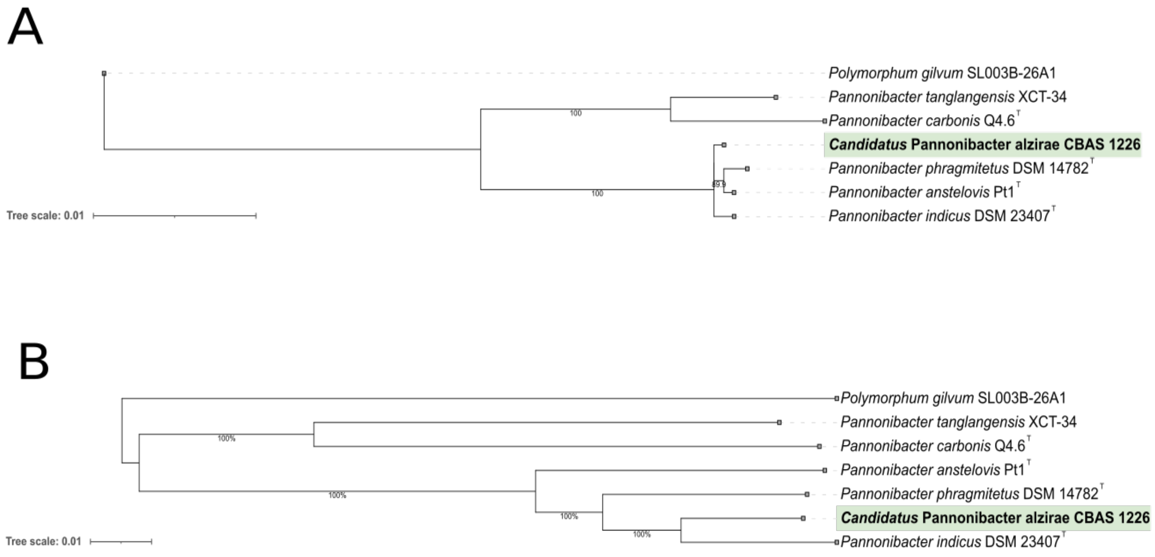


**Supplementary Figure 4. Genome-based phylogenetic relationships among *Pannonibacter* type strains, inferred using two independent methods.** (**A**) Maximum-likelihood tree inferred with IQ-TREE 2 from the concatenated alignment of 119 of the 120 bacterial single-copy marker genes used by GTDB-Tk. The alignment, comprising 5,010 masked amino acid positions, was generated using the GTDB-Tk v2.7.1 identify and align pipeline. The tree was inferred under the LG+F+G substitution model. Node support was assessed using the SH-like approximate likelihood ratio test (SH-aLRT) with 1,000 replicates. Branch lengths represent amino acid substitutions per site. (**B**) Whole-genome distance-based tree inferred using the GBDP approach. Pairwise intergenomic distances were calculated using the "trimming" algorithm and distance formula d5, with 100 distance replicates. A balanced minimum-evolution tree was inferred using FastME v2.1.6.1 with subtree pruning and regrafting (SPR) post-processing. Branch support values represent GBDP pseudo-bootstrap support estimated from 100 replicates. Branch lengths represent GBDP distances calculated using formula d5. Both trees include the type strains of all published *Pannonibacter* species, and the novel strain *Candidatus* P. alzirae CBAS1226 described in this study. *Polymorphum gilvum* SL003B-26A1 was used as the outgroup. Both trees were rooted with the same outgroup and are drawn to scale, indicating substitutions per site. Digital DDH-based species clustering performed by TYGS (70% dDDH radius around each type strain) assigned each of the seven type strain genomes analyzed to a distinct species cluster, including *Candidatus* P. alzirae CBAS1226, providing independent quantitative support for its status as a novel species.

#### 3 SUPPLEMENTARY TABLES

**Supplementary Table S1:** Genome characteristics of *Candidatus* Pannonibacter alzirae and related *Pannonibacter* strains included in this study.

| Species | ID | Accession | Size Mb | Contigs (n) | GC (%) | CDS | CheckM2 Completeness (%) | CheckM2 Contamination (%) | 16s gene coordinate |
|---|---|---|---|---|---|---|---|---|---|
| ***Candidatus* Pannonibacter alzirae** | | | | | | | | | |
| *Candidatus* P. alzirae | **CBAS 1226** | GCF_055685825.1 | 5.5 | 38 | 63.0 | 4967 | **100.00** | 0.85 | JBVOUO010000026.1 |
| *Candidatus* P. alzirae | 00954 | GCF_003703925.1 | 5.6 | 10 | 63.0 | 5069 | 100.00 | 0.70 | QXER01000015.1 |
| *Candidatus* P. alzirae | 21H_023.bin.49 ** | GCA_046291305.1 | 3.5 | 1264 | 64.0 | 4051 | 67.88 | 2.54 | - |
| *Candidatus* P. alzirae | 31801 | GCF_001484065.1 | 5.7 | 2 | 63.0 | 5131 | 100.00 | 1.17 | CP013068.1 |
| *Candidatus* P. alzirae | BB | GCF_003574985.1 | 5.5 | 3 | 63.0 | 4967 | 100.00 | 0.57 | CP032312.1 |
| *Candidatus* P. alzirae | CGMCC9175 | GCF_001188555.1 | 5.6 | 40 | 64.0 | 5084 | 100.00 | 1.18 | LGSQ01000004.1 |
| *Candidatus* P. alzirae | CS1GBFMT1 | GCF_900185695.1 | 5.2 | 31 | 63.0 | 4660 | 99.99 | 0.34 | FYAP01000025.1 |
| *Candidatus* P. alzirae | EBPR_s431bin.16 | GCF_041917155.1 | 5.3 | 70 | 64.0 | 4801 | 100.00 | 0.63 | JAYRYE010000005.1 |
| *Candidatus* P. alzirae | H1-4 | GCF_003723955.1 | 5.4 | 3 | 64.0 | 4926 | 100.00 | 0.53 | CM011124.1 |
| *Candidatus* P. alzirae | H1_Bin_Bin_32_1 ** | GCA_937928815.1 | 4.0 | 436 | 64.0 | 3828 | 83.28 | 0.27 | - |
| *Candidatus* P. alzirae | I15F10I1 | GCF_900185665.1 | 5.0 | 29 | 63.0 | 4516 | 99.99 | 0.71 | FYAN01000025.1 |
| *Candidatus* P. alzirae | P2PFMT1 | GCF_900185675.1 | 5.1 | 30 | 63.0 | 4620 | 99.99 | 0.36 | FYAO01000024.1 |
| *Candidatus* P. alzirae | R1_Bin_Bin_95_sub_1 ** | GCA_937997675.1 | 4.0 | 421 | 64.0 | 3832 | 78.58 | 0.04 | - |
| *Candidatus* P. alzirae | TSB10GB1 | GCF_900185705.1 | 5.2 | 30 | 63.0 | 4660 | 99.99 | 0.34 | FYAQ01000018.1 |
| **Other *Pannonibacter* species** | | | | | | | | | |

| | | | | | | | | | |
|---|---|---|---|---|---|---|---|---|---|
| *Pannonibacter indicus* | 21H_022.bin.4 | GCF_046296595.1 | 4.4 | 13 | 64.0 | 4063 | 99.99 | 0.00 | - |
| *Pannonibacter sp.* | 21I_055.bin.29 ** | GCA_046314515.1 | 4.6 | 195 | 64.0 | 4215 | 87.58 | 1.06 | - |
| *Pannonibacter sp.* | 21I_057.bin.14 ** | GCA_046296015.1 | 4.3 | 75 | 63.0 | 3960 | 87.52 | 0.13 | - |
| *Pannonibacter sp.* | 905_3 | GCF_050510895.1 | 4.6 | 2 | 62.0 | 4729 | 93.77 | 0.34 | JBNINT010000001.1 |
| *Pannonibacter phragmitetus* | DSM 14782$^T$ | GCF_000382365.1 | 4.8 | 40 | 63.0 | 4241 | 99.99 | 0.02 | KB908247.1 |
| *Pannonibacter indicus* | DSM 23407$^T$ | GCF_001418225.1 | 4.2 | 36 | 63.0 | 3844 | 99.99 | 0.00 | LIPT01000031.1 |
| *Pannonibacter anstelovis* | Pt1$^T$ | GCF_036881715.1 | 4.6 | 45 | 63.0 | 4188 | 99.99 | 0.10 | JBAKBF010000025.1 |
| *Pannonibacter anstelovis* | Pt2 | GCF_036881675.1 | 4.6 | 36 | 63.0 | 4179 | 99.99 | 0.10 | JBAKBE010000020.1 |
| *Pannonibacter carbonis* | Q4.6$^T$ | GCF_003012935.1 | 4.8 | 33 | 64.0 | 4359 | 100.00 | 0.10 | PUDR01000029.1 |
| *Pannonibacter indicus* | S1_Bin_MAXBIN_004_1_1 ** | GCA_938030715.1 | 4.0 | 24 | 63.0 | 3673 | 86.62 | 1.14 | - |
| *Pannonibacter sp.* | SL95 | GCF_026672425.1 | 5.0 | 6 | 64.0 | 4750 | 100.00 | 0.70 | JAPPRC010000004.1 |
| *Pannonibacter tanglangensis* | XCT-34$^T$ | GCF_009915685.1 | 4.6 | 26 | 67.0 | 4091 | 100.00 | 0.20 | JAABLP010000012.1 |
| *Pannonibacter tanglangensis* | XCT-53 | GCF_009915765.1 | 4.4 | 7 | 67.0 | 3936 | 100.00 | 0.25 | JAABLQ010000006.1 |
| **Outgroups** | | | | | | | | | |
| *Agrobacterium tomkonis* | CFBP 6623$^T$ | GCF_900013535.1 | 5.5 | 4 | 59.0 | 5171 | 100.00 | 0.51 | LT009724.1 |
| *Polymorphum gilvum* | SL003B-26A1$^T$ | GCF_000192745.1 | 4.7 | 2 | 67.0 | 4426 | 100.00 | 0.31 | CP002568.1 |

Legend: RefSeq: reference sequence accession number; ID: bacterial designation; Size (Mb): genome size in megabases; Contigs (n): number of contigs in the assembly; GC (%): genomic GC content percentage; CDS: number of predicted coding sequences; CheckM2 Completeness (%): genome completeness estimated using CheckM2; CheckM2 Contamination (%): genome contamination estimated using CheckM2.

**Supplementary Table S2: Ribosomal Multilocus Sequence Typing (rMLST) scores**

| Species | ID | Accession | Size Mb | Contigs (n) | GC (%) | CDS | rMLST (53 genes) |
|---|---|---|---|---|---|---|---|
| ***Candidatus* Pannonibacter alzirae** | | | | | | | |
| *Candidatus* P. alzirae | **CBAS 1226** | GCF_055685825.1 | 5.5 | 38 | 63.0 | 4967 | 26% *P. phragmitetus*/ 18% *P. indicus* |
| *Candidatus* P. alzirae | 954 | GCF_003703925.1 | 5.6 | 10 | 63.0 | 5069 | 36% *P. phragmitetus* |
| *Candidatus* P. alzirae | 21H_023.bin.49 ** | GCA_046291305.1 | 3.5 | 1264 | 64.0 | 4051 | 35% *P. indicus* / 35% *P. phragmitetus* |
| *Candidatus* P. alzirae | 31801 | GCF_001484065.1 | 5.7 | 2 | 63.0 | 5131 | 33% *P. phragmitetus* |
| *Candidatus* P. alzirae | BB | GCF_003574985.1 | 5.5 | 3 | 63.0 | 4967 | 28% *P. phragmitetus* |
| *Candidatus* P. alzirae | CGMCC9175 | GCF_001188555.1 | 5.6 | 40 | 64.0 | 5084 | 33% *P. phragmitetus* |
| *Candidatus* P. alzirae | CS1GBFMT1 | GCF_900185695.1 | 5.2 | 31 | 63.0 | 4660 | 28% *P. indicus* / 71% *P. phragmitetus* |
| *Candidatus* P. alzirae | EBPR_s431bin.16 | GCF_041917155.1 | 5.3 | 70 | 64.0 | 4801 | 17% *P. indicus* / 23% *P. phragmitetus* |
| *Candidatus* P. alzirae | H1-4 | GCF_003723955.1 | 5.4 | 3 | 64.0 | 4926 | 33% *P. phragmitetus* |
| *Candidatus* P. alzirae | H1_Bin_Bin_32_1 ** | GCA_937928815.1 | 4.0 | 436 | 64.0 | 3828 | 14% *P. indicus* / 21% *P. phragmitetus* |
| *Candidatus* P. alzirae | I15F10I1 | GCF_900185665.1 | 5.0 | 29 | 63.0 | 4516 | 25% *P. indicus* / 75% *P. phragmitetus* |
| *Candidatus* P. alzirae | P2PFMT1 | GCF_900185675.1 | 5.1 | 30 | 63.0 | 4620 | 28% *P. indicus* / 71% *P. phragmitetus* |
| *Candidatus* P. alzirae | R1_Bin_Bin_95_sub_1 ** | GCA_937997675.1 | 4.0 | 421 | 64.0 | 3832 | 23% *P. indicus* / 11% *P. phragmitetus* |
| *Candidatus* P. alzirae | TSB10GB1 | GCF_900185705.1 | 5.2 | 30 | 63.0 | 4660 | 28% *P. indicus* / 71% *P. phragmitetus* |
| **Other *Pannonibacter* species** | | | | | | | |
| *Pannonibacter indicus* | 21H_022.bin.4 | GCF_046296595.1 | 4.4 | 13 | 64.0 | 4063 | 58% *P. indicus* |
| *Pannonibacter sp.* | 21I_055.bin.29 ** | GCA_046314515.1 | 4.6 | 195 | 64.0 | 4215 | 66% *P. carbonis* / 33% *P. tanglangensis* |

| | | | | | | | |
|---|---|---|---|---|---|---|---|
| *Pannonibacter sp.* | 21I_057.bin.14 ** | GCA_046296015.1 | 4.3 | 75 | 63.0 | 3960 | 100% *P. carbonis* |
| *Pannonibacter sp.* | 905_3 | GCF_050510895.1 | 4.6 | 2 | 62.0 | 4729 | 6% *P. anstelovis* |
| *Pannonibacter phragmitetus* | DSM 14782$^T$ | GCF_000382365.1 | 4.8 | 40 | 63.0 | 4241 | 100% *P. phragmitetus* |
| *Pannonibacter indicus* | DSM 23407$^T$ | GCF_001418225.1 | 4.2 | 36 | 63.0 | 3844 | 100% *P. indicus* |
| *Pannonibacter anstelovis* | Pt1$^T$ | GCF_036881715.1 | 4.6 | 45 | 63.0 | 4188 | 100% *P. anstelovis* |
| *Pannonibacter anstelovis* | Pt2 | GCF_036881675.1 | 4.6 | 36 | 63.0 | 4179 | 100% *P. anstelovis* |
| *Pannonibacter carbonis* | Q4.6$^T$ | GCF_003012935.1 | 4.8 | 33 | 64.0 | 4359 | 100% *P. carbonis* |
| *Pannonibacter indicus* | S1_Bin_MAXBIN_004_1_1 ** | GCA_938030715.1 | 4.0 | 24 | 63.0 | 3673 | 94% *P. indicus* |
| *Pannonibacter sp.* | SL95 | GCF_026672425.1 | 5.0 | 6 | 64.0 | 4750 | 66% *P. carbonis* / 33% *P. tanglangensis* |
| *Pannonibacter tanglangensis* | XCT-34$^T$ | GCF_009915685.1 | 4.6 | 26 | 67.0 | 4091 | 100% *P. tanglangensis* |
| *Pannonibacter tanglangensis* | XCT-53 | GCF_009915765.1 | 4.4 | 7 | 67.0 | 3936 | 100% *P. tanglangensis* |
| **Outgroups** | | | | | | | |
| *Agrobacterium tomkonis* | CFBP 6623$^T$ | GCF_900013535.1 | 5.5 | 4 | 59.0 | 5171 | 100% *A. tomkonis* |
| *Polymorphum gilvum* | SL003B-26A1$^T$ | GCF_000192745.1 | 4.7 | 2 | 67.0 | 4426 | 100% *P. gilvum* |

Legend: ID: bacterial designation; Size (bp): genome size in base pairs; Contigs (n): number of contigs in the assembly; GC (%): genomic GC content percentage; CDS (Prokka): number of coding sequences predicted by Prokka; rMLST (53 genes): number of ribosomal genes identified for sequence typing. Genomes indicated by asterisks (**) were recovered from metagenomic datasets (MAGs); consequently, their assembly size and gene content may be underestimated due to incomplete genome reconstruction.

**Supplementary Table S3: Biochemical characterization of *Candidatus* Pannonibacter alzirae and comparison with related species, with comparative phenotypic data for reference strains compiled from the literature and databases.**

| Biochemical test | *Candidatus* Pannonibacter alzirae | *Pannonibacter phragmitetus* DSM 14782 | *Pannonibacter indicus* DSM 23407 |
|---|---|---|---|
| 1-O-methyl-alpha-D-glucoside as carbon source | - | - | ND |
| 2-ketoglutarate as carbon source (2KT) | + | - | ND |
| 4-Methylumbelliferyl | - | ND | ND |
| 5,5'-dithiobis(2-nitrobenzoic acid) | + | ND | ND |
| 5-ketogluconate as carbon source (5KG) | - | + | ND |
| Acetate | + | ND | ND |
| Acid production from d-glucose | - | - | + |
| Alanine-Phenylalanine-Proline-Arylamidase | - | ND | ND |
| Alkaline phosphatase | - | + | + |
| Alkalinization of L-Lactate | + | ND | ND |
| α-Galactosidase | + | + | + |
| α-Glucosidase | + | + | - |
| Arginine-arginine | - | ND | ND |
| β-D-Alose | - | ND | ND |
| β-Galactosidase | + | + | + |
| β-Glucosidase | + | - | - |
| β-Glucuronidase | - | - | - |
| β-Alanine-Arylamidase | - | ND | ND |
| β-Xylosidase | - | ND | ND |
| B-Gentiobiose | - | ND | ND |
| Bisphosphate | + | ND | ND |
| Cinnamate | - | ND | ND |
| Colistin | + | ND | ND |
| D-Arabitol as carbon source (DARL) | - | - | + |
| D-Cellobiose as carbon source (CEL) | - | + | + |
| Dextrose | - | ND | ND |
| D-Fructose as carbon source (FRU) | - | + | + |
| D-Galactose as carbon source (GAL) | - | + | ND |
| D-Galacturonate as carbon source (GAT) | - | + | ND |

| | | | |
|---|---|---|---|
| D-Glucose as carbon source (GLU) | - | + | + |
| D-Mannose as carbon source (MNE) | - | + | + |
| D-Melezitose as carbon source (MLZ) | - | - | ND |
| D-Melibiose as carbon source (MEL) | - | + | + |
| D-Sorbitol as carbon source (SOR) | - | - | ND |
| D-Tagatose as carbon source (TAG) | - | - | ND |
| D-Trehalose as carbon source (TRE) | - | + | ND |
| Esculin as carbon source (ESC) | - | + | + |
| γ-Glutamyl-Transferase | - | ND | ND |
| Gentiobiose as carbon source (GEN) | - | + | ND |
| Gluconic acid | - | ND | ND |
| Glutamyl-Arylamidase | - | ND | ND |
| Glutamyl-Glycyl-Arginine-Arylamidase | - | ND | ND |
| Glycine | + | ND | ND |
| Glycine Arylamidase | - | ND | ND |
| Glycine-proline | - | ND | ND |
| H2S production | - | - | - |
| L-Arabinose as carbon source (ARA) | - | + | ND |
| L-Arginine | - | + | + |
| Leucine arylamidase | + | + | + |
| L-γ-glutamyl-p-nitroanilide | - | ND | ND |
| L-Glutamate as carbon source (GTT) | + | + | ND |
| L-Histidine as carbon source (HIS) | - | - | ND |
| Lipase (C 14) | - | - | - |
| L-Lactate Assimilation | - | ND | ND |
| L-Malate as carbon source (LMLT) | - | - | ND |
| L-Phenylalanine | - | ND | - |
| L-Proline Arylamidase | + | ND | ND |
| L-Proline as carbon source (PRO) | + | - | ND |
| L-Proline-p-nitroalanilide | + | ND | ND |
| L-Pyroglutamic acid | + | + | ND |
| L-Pyrrolidonyl Arylamidase | + | ND | ND |
| L-Rhamnose | - | + | ND |

| | | | |
|---|---|---|---|
| L-Rhamnose as carbon source (RHA) | - | + | ND |
| L-Tryptophan | - | ND | ND |
| Lysine decarboxylase | - | ND | + |
| Lysine-alanine | - | ND | ND |
| Maltose as carbon source (MAL) | - | + | ND |
| Maltulose | - | ND | ND |
| N-Acetyl-β-glucosaminidase | - | - | - |
| N-Acetyl-D-glucosamine as carbon source (NAG) | - | - | ND |
| N-Acetyl-galactosamine | - | ND | ND |
| N-Glutaryl-glycine-arginine | - | ND | ND |
| Ornithine | - | ND | - |
| Palatinose as carbon source (PLE) | - | + | ND |
| P-Nitrophenyl-B-D-glucoside | + | ND | ND |
| Polymyxin B | + | ND | ND |
| Resistance to O/129 (*Vibrio* compound) | - | ND | ND |
| Succinate as carbon source (SUC) | + | + | ND |
| Sucrose as carbon source (SAC) | - | + | ND |
| Tiglic acid | + | ND | ND |
| Tryptophan as carbon source (TRY) | - | - | ND |
| Tyrosine Arylamidase | + | ND | ND |

+: Positive; -: Negative; ND: Data not provided.

**Supplementary Table S4: Minimum inhibitory concentrations of *Candidatus* Pannonibacter alzirae CBAS 1226 to evaluated antimicrobials**

| **Antimicrobial** | **MIC (μg mL$^{-1}$)** |
|---|---|
| Amikacin | 8 |
| Amoxicillin-Clavulanic acid | >32 |
| Ampicillin-Sulbactam | >32 |
| Aztreonam | >64 |
| Cephalexin | ≤4 |
| Cefepime | 16 |
| Ceftazidime-Avibactam | 4 |
| Ceftazidime | 16 |
| Ceftolozane-Tazobactam | 16 |
| Ceftriaxone | 1 |
| Cefuroxime | ≤1 |
| Cefuroxime-Axetil | ≤1 |
| Ciprofloxacin | ≤0.06 |
| Ertapenem | ≤0.12 |
| Fosfomycin | >256 |
| Gentamicin | 2 |
| Imipenem | < 0.25 |
| Meropenem | < 0.25 |
| Nitrofurantoin | <16 |
| Norfloxacin | < 0.5 |
| Piperacillin-Tazobactam | ≥128 |
| Polymyxin B | >256 |
| Sulfamethoxazole-Trimethoprim | 0.06 |
| Tigecycline | < 0.5 |

**MIC** - Minimum Inhibitory Concentration.

**Supplementary Table S5: Identified protein domains associated with antimicrobial resistance in *Candidatus* Pannonibacter alzirae CBAS 1226**

| Domain description | Antimicrobial putative interaction | InterProScan accession |
|---|---|---|
| Aminoglycoside phosphotransferase | Aminoglycoside | IPR002575 |
| AmpG-like permease/Acetyl-coenzyme A transporter 1 | Penicillin, Cephamycin, Cephalosporin | IPR004752 |
| Beta-lactamase, class-A active site | Penicillin, Cephamycin, Cephalosporin | IPR023650 |
| Beta-lactamase/transpeptidase-like | Cefepime, Cefotaxime, Ceftazidime, Piperacillin | IPR012338 |
| Beta-lactamase-related | Penicillin, Cephamycin, Cephalosporin | IPR001466 |
| Chloramphenicol acetyltransferase-like domain superfamily | Chloramphenicol | IPR023213 |
| Drug resistance transporter Bcr/CmlA subfamily | Bicyclomycin, Chloramphenicol, Florfenicol | IPR004812 |
| Drug resistance transporter EmrB-like | Multidrug | IPR004638 |
| Glyoxalase/fosfomycin resistance/dioxygenase domain | Fosfomycin | IPR004360 |
| Multi antimicrobial extrusion protein | Multidrug | IPR002528 |
| Multidrug efflux transporter AcrB TolC docking domain | Multidrug | IPR027463 |
| Multiple antibiotic resistance (MarC)-related | Multidrug | IPR002771 |
| PBP domain | Penicillin | IPR024370 |
| Penicillin binding protein transglycosylase domain | Cefepime, Cefotaxime, Ceftazidime, Piperacillin | IPR036950 |
| Penicillin-binding protein, C-terminal domain superfamily | Piperacillin | IPR015956 |

| | | |
|---|---|---|
| Penicillin-binding protein, dimerisation domain | Cefepime, Cefotaxime, Ceftazidime, Piperacillin | IPR005311 |
| Penicillin-binding protein, OB-like domain | Cefepime, Ceftazidime | IPR031376 |
| Penicillin-binding protein, transpeptidase | Cefepime, Cefotaxime, Ceftazidime, Piperacillin | IPR001460 |
| Penicillin-binding, C-terminal | Penicillin | IPR009647 |
| Peptidase M74, penicillin-insensitive murein endopeptidase | Penicillin | IPR005073 |
| PsrA, tetracycline repressor-like, C-terminal domain | Tetracycline | IPR041586 |
| Small drug/metabolite transporter protein family | Multidrug | IPR000390 |
| Small multidrug resistance protein | Multidrug | IPR045324 |
| Tetracycline repressor-like, C-terminal domain | Tetracycline | IPR011075 |
| Tetracycline repressor-like, C-terminal domain superfamily | Tetracycline | IPR036271 |
| Tetracycline repressor TetR, C-terminal | Tetracycline | IPR004111 |
| Tetracycline resistance protein TetA/multidrug resistance protein MdtG | Tetracycline | IPR001958 |

**Supplementary Table S6. Pairwise genomic similarity between strain CBAS 1226 and representative *Pannonibacter* genomes included in this study**

| Genome | Current NCBI annotation | ANI vs. CBAS 1226 (%) | AAI vs. CBAS 1226 (%) | Phylogenomic clustering | Proposed assignment |
|---|---|---|---|---|---|
| ***Candidatus*_P_alzirae_*EBPR_s431bin.16*** | ***Pannonibacter sp.*** | **99.69** | **99.79** | **Same lineage** | ***Candidatus* P. alzirae** |
| ***Candidatus*_P_alzirae_*00954*** | ***P. phragmitetus*** | **97.76** | **98.52** | **Same lineage** | ***Candidatus* P. alzirae** |
| ***Candidatus*_P_alzirae_*31801*** | ***P. phragmitetus*** | **97.62** | **98.30** | **Same lineage** | ***Candidatus* P. alzirae** |
| ***Candidatus*_P_alzirae_*CGMCC9175*** | ***P. phragmitetus*** | **97.58** | **98.50** | **Same lineage** | ***Candidatus* P. alzirae** |
| ***Candidatus*_P_alzirae_*21H_023.bin.49*** | ***P. indicus*** | **97.17** | **98.29** | **Same lineage** | ***Candidatus* P. alzirae** |
| ***Pannonibacter_indicus_DSM_23407*** | ***P. indicus*$^{T}$** | **95.66** | **97.14** | **Distinct lineage** | ***P. indicus*** |
| ***Pannonibacter_indicus_S1_Bin_MAXBIN_004_1_1*** | ***P. indicus*** | **95.61** | **97.04** | **Distinct lineage** | ***P. indicus*** |
| ***P_indicus_21H_022.bin.4*** | ***P. indicus*** | **95.54** | **97.06** | **Distinct lineage** | ***P. indicus*** |
| ***Pannonibacter_phragmitetus_DSM_14782*** | ***P. phragmitetus*$^{T}$** | **94.26** | **96.11** | **Distinct lineage** | ***P. phragmitetus*** |
| ***Pannonibacter_anstelovis_Pt1*** | ***P. anstelovis*$^{T}$** | **91.91** | **94.75** | **Distinct lineage** | ***P. anstelovis*** |
| ***Pannonibacter_anstelovis_Pt2*** | ***P. anstelovis*** | **91.89** | **94.77** | **Distinct lineage** | ***P. anstelovis*** |
| ***Pannonibacter_tanglangensis_XCT-34*** | ***P. tanglangensis*$^{T}$** | **81.43** | **76.01** | **Distinct lineage** | ***P. tanglangensis*** |
| ***Pannonibacter_XCT-53*** | ***P. tanglangensis*** | **81.38** | **76.09** | **Distinct lineage** | ***P. tanglangensis*** |
| ***Pannonibacter_carbonis_Q4.6*** | ***P. carbonis*** | **81.14** | **75.98** | **Distinct lineage** | ***P. carbonis*** |

Legend: Genomes clustering with *Candidatus* P. alzirae CBAS 1226 in phylogenomic analyses are proposed to represent members of *Candidatus* Pannonibacter alzirae sp. nov., whereas type strains and representatives of the remaining recognized species are shown for comparison. ANI and AAI values correspond to pairwise comparisons against strain CBAS 1226, calculated with FastANI and EzAAI, respectively.

**Supplementary Table S7. Functional categorization of the 537 genes exclusively found in *Candidatus* P. alzirae CBAS 1226, when compared to *P. indicus* and *P. phragmitetus* type strains.**

| Functional category | No. of genes | Biological interpretation |
|---|---|---|
| Type IV secretion system (T4SS) | 5 | Conjugation machinery and horizontal DNA transfer |
| CRISPR-Cas (Type I) | 4 | Adaptive immunity against bacteriophages and exogenous DNA |
| Restriction-modification (R-M) system | 3 | Innate defense against invading DNA |
| Toxin-antitoxin (TA) systems | 5 | Stabilization of mobile genetic elements and persistence |
| Antimicrobial/antiseptic resistance | ~9 | Efflux systems, β-lactam hydrolysis, and quaternary ammonium compound (QAC) resistance |
| Metal resistance/homeostasis | ~6 | Adaptation to Cd, Co, Zn, chromate, and copper stress |
| Hydrogen metabolism (NiFe hydrogenase) | 7 | Potential utilization of $H_2$ as an auxiliary energy source |
| Heme/iron acquisition | 3 | Iron acquisition under nutrient-limited conditions |
| Quorum sensing/quorum quenching and biofilm formation | 5 | Modulation of surface-associated behavior and interspecies competition |

| | | |
|---|---|---|
| Transcriptional regulators | 46 | Expanded regulatory and adaptive capacity |
| Hypothetical proteins with no functional annotation | 270 | 50.3% of the total; candidates for future functional characterization |

**Supplementary Table S8. Comparative taxonomic descriptions of previously published Pannonibacter species.**

| **Species** | **Etymology** | **Type strain** | **Isolation source / origin** | **Cell morphology and motility** | **Colony morphology** | **Growth range (temperature / pH / NaCl)** | **Oxygen requirement** | **Catalase / oxidase** | **Selected biochemical characteristics** | **DNA G+C / genome size** | **Reference** |
|---|---|---|---|---|---|---|---|---|---|---|---|
| *Pannonibacter phragmitetus* | N.L. masc. adj. *phragmitetus*, of the plant association Scirpo-Phragmitetum, referring to the habitat. | DSM 14782$^T$ | Surface of decomposing reed (Phragmites australis) rhizomes, Lake Fertő, Hungary. | Gram-stain-negative, non-spore-forming, straight to slightly curved rods; motile by polar flagella; approximately 2.0–4.0 × 0.3–0.6 | On alkaline agar: whitish-beige, circular, convex, smooth, shiny, entire margins. | 15–45 °C; pH 7.0–11.0; 0–5% NaCl. Optimum temperature reported around 30–37 °C. | Facultatively anaerobic; chemo-organotrophic. | Positive / positive. | Acid from D-glucose under aerobic and anaerobic conditions; nitrate reduced to nitrogen; arginine hydrolysis, citrate utilization, urease and phosphatase positive; aesculin, casein, cellulose, gelatin, hippurate, starch and Tween 80 hydrolysis negative; indole, H2S, MR and VP negative; BChl a not synthesized aerobically. | 64.6 mol% G+C in the original description; approximately 4.78 Mbp genome and 63.1% genomic G+C in the later type-strain genome analysis. | Borsodi et al. 2003, Int J Syst Evol Microbiol 53:555–561; Hördt et al. 2020, Front Microbiol 11:468. |

| | | | | | | | | | | | |
|---|---|---|---|---|---|---|---|---|---|---|---|
| | | | | µm; PHA-containing. | | | | | | | |
| *Pannonibacter indicus* | Corrected name; originally published as *Pannonibacter indica*. N.L. masc. adj. indicus, of India. | DSM 23407$^{T}$ | Hot-spring sediment, Athamallik, Odisha (Orissa), India. | Gram-negative, motile rods; strictly aerobic. | On nutrient agar: smooth, round, cream-coloured. | 20–45 °C; pH 6.0–11.0; reported optimum 37 °C and pH 7.0–10.0. NaCl tolerance up to approximately 7% in comparative testing. | Strictly aerobic. | Positive / positive. | Urease, phosphatase, growth on MacConkey, King's A and King's B, lysine decarboxylase, aesculin hydrolysis, β-galactosidase, citrate, casein and arginine hydrolysis positive. Nitrate/nitrite reduction, MR, VP, indole, phenylalanine deamination, malonate, ornithine decarboxylase, pigment, DNase, H2S, cetrimide growth, starch/cellulose/gelatin/hippurate hydrolysis and lecithinase negative. Strong arsenate tolerance. | 63.3 mol% G+C for the type strain. | Bandyopadhyay et al. 2013, Arch Microbiol 195:1–8; corrected nomenclature recorded by LPSN. |
| *Pannonibacter carbonis* | L. gen. masc. n. *carbonis*, of coal/charcoal. | Q4.6$^{T}$ | Coal-mine water from a coal-producing well in Jincheng, Shanxi | Gram-stain-negative, rod-shaped, motile by a single polar flagellum | On TSA: smooth, cream-coloured, circular; approximately 1–1.6 mm diameter. | 4–55 °C (optimum 30–35 °C); pH 3.5–10.5 (optimum 6.5–8.0); 0–5.5% NaCl | Facultatively anaerobic. | Positive / positive. | Urease, VP, gelatin hydrolysis and H2S positive; nitrate reduction, indole and Tween 80 hydrolysis negative. Utilizes D-galactose, D-glucose, maltose, trehalose, ONPG, arginine, | 63.6 mol% G+C; genome approximately 4.78 Mbp for Q4.6T. | Xi et al. 2018, Int J Syst Evol Microbiol 68:2042–2047. |

| | | | | | | | | | | | |
|---|---|---|---|---|---|---|---|---|---|---|---|
| | | | Province, China. | ; 1.8–3.0 × 0.3–0.7 µm; PHA-containing. | | (optimum 0–2%). | | | lysine, ornithine, citrate, sorbitol, mannitol, potassium gluconate, xylose and xylitol; sucrose weakly utilized. Does not utilize arabinose, erythritol, mannose, lactose, adipic acid, inositol or starch. | | |
| *Pannonibacter anstelovis* | Name proposed by Castellana et al.; the article presents it as a novel/putative species associated with bloodstream infections. | Pt1$^{T}$ (reference genome) | Blood cultures from two paediatric patients in a hospital paediatric oncohaematology unit, Bari, Italy. | Gram-negative bacilli/rod-shaped cells; detailed cell dimensions and flagellation were not reported in the species proposal. | Milky-white, mucous colonies with an acrid odour on PVX chocolate agar after 24 h at 37 °C in 5% CO2. | Growth conditions were not characterized as comprehensively as in the formal species descriptions; draft genomes were approximately 4.6 Mbp and GC content was 62.58% (62.78% in the ONT assembly). | Growth demonstrated under microaerophilic/5% CO2 conditions; full aerobic/anaerobic physiological characterization not reported. | Not comprehensively reported. | API 20E: ONPG and citrate positive; ADH, LDC, ODC, H2S, urease, TDA, indole, VP, gelatin and nitrate negative. High MICs to aminoglycosides, third-/fourth-generation cephalosporins and piperacillin/tazobactam; susceptible to ciprofloxacin and carbapenems. | 62.58% genomic G+C; approximately 4.6 Mbp draft genomes. | Castellana et al. 2024, Microorganisms 12:799. DOI: 10.3390/microorganisms12040799. Note: LPSN currently lists the name as not validly published under the ICNP. |

| | | | | | | | | | | | |
|---|---|---|---|---|---|---|---|---|---|---|---|
| *Pannonibacter tanglangensis* | N.L. masc. adj. *tanglangensis*, referring to Tanglang Mountain, Shenzhen, China, where the species was isolated. | XCT-34T = XCT-34$^{T}$ | Sediment from an artificial freshwater reservoir near Tanglang Mountain, Shenzhen, China. | Aerobic, Gram-stain-negative, rod-shaped; motile by polar flagella; approximately 1.4–3.4 × 0.4–0.9 µm. | After 2 days on R2A at 35 °C: ivory, circular, smooth, entire margin. | 10–40 °C (optimum 35–37 °C); pH 4.0–10.0 (optimum pH 8.0); up to 4% NaCl (optimum 0.5%). | Strictly aerobic. | Positive / negative. | API 50CH: acid from D-fructose, D-fucose, D-glucose, D-mannose, D-xylose, L-fucose, maltose, polychrome esculin citrate, sucrose and trehalose; weakly from D-arabinose, D-galactose, D-ribose, D-turanose, cellobiose, glycerol, inositol, L-arabinose, melibiose, N-acetylglucosamine and potassium 5-ketogluconate. API ZYM positive for acid/alkaline phosphatases, esterases, leucine arylamidase, naphthol-AS-BI-phosphohydrolase, phosphohydrolase, trypsin, valine arylamidase, α-galactosidase, β-galactosidase, β-glucosidase and β-glucuronidase. | 67.2% genomic G+C; circular chromosome 4,588,697 bp. | Wang et al. 2024, Journal of Microbiology 62:727–737. DOI: 10.1007/s12275-024-00151-y; validly published in Validation List no. 223 (2025). |
